# Cholesterol Dysregulation in APOE4 Astrocytes Promotes α-Synuclein Pathology in miBrains, a Human Brain Tissue Model

**DOI:** 10.1101/2025.02.09.637107

**Authors:** Louise A. Mesentier-Louro, Camille Goldman, Sebastian Gaese, Alice Buonfiglioli, Dimitrios Kyriakis, Ashley Harlock, Alain Ndayisaba, Emily R. Sartori, Abigail Uchitelev, John F. Fullard, Evelyn Hennigan, Donghoon Lee, Braxton R. Schuldt, Rikki B. Rooklin, Jonathan Barra, Jose Javier Bravo-Cordero, Panos Roussos, Vikram Khurana, Joel W. Blanchard

## Abstract

The pathological hallmarks of neurodegeneration are the aberrant post-translational modification and aggregation of proteins. Genetic factors, like *APOE4,* increase the prevalence and severity of tau, amyloid, and α-synuclein pathologies. However, the human brain is largely inaccessible during this process, limiting mechanistic understanding. Here, we developed an iPSC-based 3D model that integrates neurons, glia, myelin, and cerebrovascular cells into a human brain-like tissue (“miBrain”). Single-nucleus RNA sequencing of miBrains confirmed the presence of diverse cell populations and revealed transcriptional responses to α-synuclein pathology. Like the human brain, pathogenic α-synuclein is increased in *APOE4/4* miBrains. Combinatorial experiments revealed that endolysosomal dysfunction caused by cholesterol accumulation in *APOE4/4* astrocytes impairs the degradation of soluble α-synuclein leading to a pathogenic transformation that seeds α-synuclein inclusions in neurons. Collectively, this study establishes a robust model for investigating protein inclusions in human iPSC-derived brain tissue and highlights the role of astrocytes and cholesterol in *APOE4*-mediated pathologies.

## Introduction

Alzheimer’s disease (AD) is canonically associated with amyloid-β and tau pathology^1^, yet aggregated α-synuclein (α-Syn) inclusions occur in 50-90% of cases^2–5^ and accelerate neurodegeneration^5–10^ and cognitive decline^11,12^. *APOE4*, the strongest genetic risk factor for AD, significantly increases α-Syn pathology in AD^12,13^ and is a major genetic risk factor for Lewy Body Dementia (LBD)^14,15^. However, how *APOE4* promotes α-Syn pathology remains unclear. Elucidating these mechanisms may uncover new therapeutic and diagnostic opportunities for AD and LBD.

Faithful models of α-Syn pathology are essential to uncover how genetic factors like *APOE4* modify neurodegeneration. Although α-Syn pre-formed fibrils (PFFs) are commonly employed to induce pathology^16–18^ in mice^19,20^ and *in vitro*^17,21,22^, these models are variable and pose biohazard concerns^23^. This highlights the need for tractable, PFF-independent systems that enable investigation of multicellular mechanisms. Traditional two-dimensional (2D) cultures may also fail to recapitulate the brain microenvironment^24^, whereas three-dimensional (3D) systems more effectively reproduce neuropathological phenotypes^25,26^.

We developed the human **m**ulti-cellular **i**ntegrated **Brain** (**miBrain**) ^27^, an induced pluripotent stem cell (iPSC)-derived 3D human brain tissue containing functional neurons, oligodendroglia, astrocytes, microglia, and a blood-brain barrier-like vasculature. As an engineered tissue, isogenic miBrains can be assembled with AD risk variants like *APOE4* in specific cell types. Here, we further advance the platform by introducing cryopreservation methods that enable batch production, improving the reproducibility and scalability of the miBrain. Using this platform, we established a highly reproducible model of α-Syn neuropathological accumulation and applied it to investigate how *APOE4* influences α-Syn pathology. Our findings identify APOE4 in astrocytes causes lipid dysregulation that impairs lysosomal function ultimately promoting α-Syn pathologies in neurons, opening new therapeutic opportunities for AD, LBD, and related diseases.

## Results

### miBrains: A human brain microphysiological system

The miBrain is a human brain-like tissue generated by encapsulation of iPSC-derived neurons^28,29^, astrocytes^30,31^, endothelial cells^32–35^, mural cells^32,36^, oligodendrocyte precursor cells (OPCs)^37^, and microglia^38^. miBrains displayed homogenous TUJ1^+^ neuronal networks and PECAM1^+^ vascular structures (**Figure 1a**). Staining for multiple markers revealed interactions and co-localization between S100b^+^ and AQP4^+^ astroglia and PECAM1^+^ vascular structures, vascular coverage with NG2^+^ mural cells, myelin basic protein (MBP)^+^/neurofilament^+^ neuro-myelin structures, and the presence of IBA1^+^ microglia (**Figure 1b**), consistent with the cell types encapsulated in the miBrain. All miBrains used in this study contained all the above-mentioned cell types, unless indicated in the figure legend.

**Figure 1.**
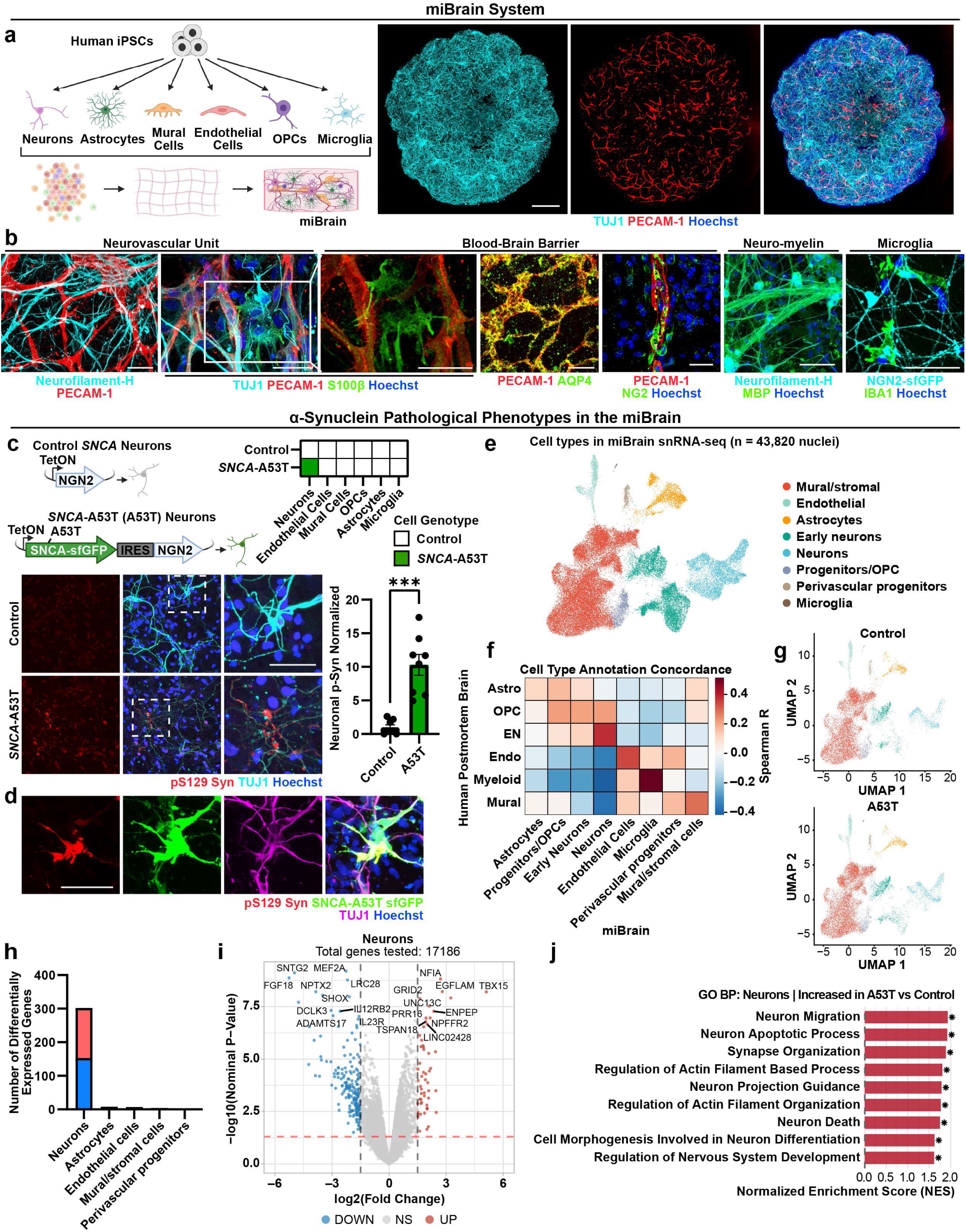
Induction of α-Syn intracellular inclusions in multi-cellular integrated Brain (miBrain) tissue. **a**. Schematic of miBrain generation and images of 4-week-old miBrains stained for TUJ1 (cyan) and PECAM-1 (red). Scale bar: 500 µm. **b**. Representative images of 2-4-week-old miBrains stained for neurons (Neurofilament-H, TUJ1, NGN2-sfGFP; cyan), astrocytes (S100b, AQP4; green), endothelial cells (PECAM-1; red), mural cells (NG2; green), myelin (MBP; green), and microglia (IBA1, CD68; green, magenta). Scale bar: 50 µm. **c.** Representative images of 18-day-old control and *SNCA*-A53T miBrains stained for α-Syn pS129 (red) and TUJ1 (cyan). α-Syn pS129 volume was measured within TUJ1-positive cells and normalized to control. Scale bar: 50 µm. Data are mean ± SEM (n = 4 biological replicates). P-value by Welch’s two-tailed t-test **d.** Representative images of α-Syn pS129 (red) inclusion co-localized with sfGFP (green) and TUJ1 (magenta) cells. Scale bar: 50 µm. **e.** UMAP of miBrain cell clusters (43,820 nuclei; Control, *APOE3/3 A53T* and *APOE4/4 A53T;* n = 2 biological replicates per condition, each containing two pooled miBrains). **f.** Correlation analysis mapping miBrain cell types to human postmortem brain dataset (T. Clarence et al)^112^. **g.** Split UMAPs showing similar cellular composition in Control and *SNCA-A53T* miBrains. **h.** Number of differentially expressed genes (DEGs) by cell type in *APOE3/3 A53T* versus control (adj. p < 0.05, |logFC| > 0.5). **i.** Volcano plot of neuronal DEGs (upregulated, red; downregulated, blue; |log2 fold change| > 1.5, p-value < 0.05). **j.** Gene Ontology (GO) biological processes enriched in *A53T APOE3/3* neurons. *p < 0.05, **p < 0.01, ***p < 0.001, ****p < 0.0001.

To functionally validate neuronal network activity within miBrains, we performed multielectrode array (MEA) recordings (**Figure S1a**). miBrains exhibited robust spontaneous electrical activity, with a mean firing rate of 1.968 Hz (± 0.171 SEM; **Figure S1b, i**) with near-complete electrode engagement (15–16 active electrodes per array; **Figure S1b, ii**). These values confirm the presence of functionally active neuronal populations in miBrains. Additionally, miBrains displayed frequent spontaneous bursts (504 ± 56 SEM; **Figure S1b, iii**) and synchronized network bursts involving an average of 6.157 electrodes (± 0.223 SEM; **Figure S1b, iv**), comparable to synchronized network activity in human brain organoids^39,40^. Together, these data suggest that miBrains recapitulate key features of developing neural networks, spatial propagation of activity across electrodes, and emergent synchronization at the network level.

In other *in vitro* platforms, such as brain organoids, significant variability in cell composition has been reported^39^. To improve miBrain reproducibility and scalability, we developed methods to cryopreserve large batches of miBrains containing multiple cell types at defined ratios (**Figure S1c**). This approach enables thawing of pre-mixed cells and addition of any desired cell type, such as neurons with genetic modifications, during tissue assembly. This eliminates the need for synchronized differentiation of multiple cell types before each experiment. Cryopreserved miBrains retained more than 90% cell viability upon thawing, expressed specific cell markers after two weeks in culture (**Figure S1c, d)**, and showed no significant differences in the ratio of neurons to nuclei across three independent batches (p = 0.292, **Figure S1d**).

### Induction of α-Syn intracellular inclusions in miBrains

Intracellular neuronal α-Syn inclusions have been challenging to reproduce *in vitro*, typically requiring long protocols^41^ and α-Syn PFFs^29^. Because phosphorylation of α-Syn on S129 is a hallmark of pathological inclusions^42,43^, we first investigated whether α-Syn PFF^17,19–22^ could induce this phenotype in miBrains. PFF inoculation significantly increased phosphorylated α-Syn (p-Syn) levels when compared to control (p = 0.0175, **Figure S1e**), demonstrating that miBrains support α-Syn pathological phenotypes. However, p-Syn appeared primarily as dispersed puncta (**Figure S1e**) rather than the neuronal inclusions characteristic of synucleinopathies^44^, and showed substantial inter-experimental variability (**Figure S1e**), consistent with previous reports^20^. Therefore, we sought a more robust approach to induce pathogenic α-Syn phenotypes in the miBrain.

A recent study found that iPSC-derived neurons express lower levels of the gene encoding α-Syn (*SNCA)* than human brain tissue and, therefore, overexpressed *SNCA* to achieve physiological levels and induce neuronal inclusions^29^. Using a similar strategy, we generated iPSC-derived neurons from a donor with familial Parkinson’s disease with inducible expression of *SNCA* bearing the A53T mutation (*SNCA-*A53T), known to enhance α-Syn’s aggregation^29,45,46^. *SNCA*-A53T was fused to superfolder green fluorescent protein (sfGFP), a variant engineered for folding and monomeric stability even when attached to aggregation-prone proteins^47–49^, thereby enabling live imaging and direct visualization of α-Syn accumulation while preserving α-Syn’s aggregation and seeding behavior^50^.

In 2D cultures, neurons with only endogenous *SNCA* expression (control) and *SNCA-*A53T neurons exhibited similar basal p-Syn levels (p = 0.959), which increased selectively in *SNCA-*A53T neurons exposed to PFFs (p< 0.0001, **Figure S1f**). This effect was absent in neurons overexpressing *SNCA*-A53T lacking the α-Syn non-amyloid component (*SNCA-*A53T-ΔNAC, p = 0.952), known to be required for α-Syn aggregation, suggesting that the observed phosphorylation arises from α-Syn biology rather than from the sfGFP fusion. In contrast to 2D cultures, miBrains with *SNCA-*A53T overexpression in neurons (*SNCA*-A53T miBrains) developed a 10.29-fold increase (±1.55 SEM) in neuronal p-Syn compared to isogenic miBrains harboring control neurons (p = 0.0004, **Figure 1c**), even without exogenous PFFs. *SNCA-*A53T neurons in 3D also displayed PFF-independent increases in p-Syn compared to control neurons (p = 0.0007; **Figure S1g)**, suggesting that the 3D environment is inherently more permissive to α-Syn pathology. PFF exposure further increased p-Syn levels in *SNCA-*A53T miBrains (p = 0.038, **Figure S1h**), consistent with seeded amplification of pathological phenotypes.

A subset of SNCA-A53T-sfGFP co-localized with p-Syn (**Figure 1d** and **Video S1**), and this fraction was similar in miBrains with or without PFFs (p = 0.275, **Figure S1i**). Most p-Syn co-localized with sfGFP, indicating phosphorylation of overexpressed SNCA-A53T (Control: 86.7% ± 1.9 SEM, PFFs: 88.0% ± 1.4 SEM). The remaining p-Syn did not co-localize with sfGFP, indicating phosphorylation of endogenous, non-A53T α-Syn in neuronal and possibly non-neuronal cells (Vehicle: 13.3% ± 1.9 SEM, PFFs: 12.0% ± 1.4 SEM**; Figure S1j**). Collectively, these results show that *SNCA*-A53T miBrains developed p-Syn-rich inclusions via corruption of both induced (A53T) and endogenous α-Syn without requiring PFFs.

Single-nucleus RNA sequencing (snRNA-seq) was performed to define the cellular composition of miBrains and to assess transcriptional changes associated with *SNCA-*A53T expression. Neuronal and glial populations including neurons, astrocytes, microglia, endothelial cells, mural cells, and OPCs were identified based on canonical marker expression (**Figure S1k**) and transcriptional similarity to a published dataset from postmortem human Parkinson’s disease brain tissue⁵¹ (**Figure 1e, f**). Quantitative analysis revealed comparable representation of major cell types between miBrains with *SNCA-*A53T neurons and miBrains with control neurons (**Figure S1l**), indicating that neuronal α-Syn pathology does not induce large-scale shifts in overall cellular composition (**Figure 1g**). These findings suggest that the model preserves multicellular architecture while enabling the assessment of cell-type-specific transcriptional responses.

Differential gene expression analysis of all cellular types showed that neurons have the highest number of differentially expressed genes (DEGs) in *SNCA-*A53T compared to control miBrains (153 upregulated, 149 downregulated, adjusted p< 0.05, |logFC| > 0.5, **Figure 1h, I, Table S1**), consistent with specific induction of *SNCA*-A53T in these cells. Gene ontology analysis revealed a significant upregulation of biological processes related to neuronal apoptosis, morphological changes, and synaptic changes (**Figure 1j, Table S2**). Gene set enrichment analysis (GSEA) further showed significant modulation of gene sets associated with α-Syn pathology (P = 0.006), synaptic vesicle maturation (P = 0.004), and protein dephosphorylation (P = 0.02; **Figure S1m**). Collectively, these transcriptional changes are consistent with neuronal dysfunction and responses to α-Syn pathogenic accumulation in *SNCA-*A53T miBrains.

### α-Syn inclusions in miBrain tissue exhibit pathological features of Lewy body-like inclusions

To assess the pathological relevance of α-Syn inclusions formed in the miBrain tissue, we examined their association with cellular features commonly observed in or near Lewy body pathology in the human brain, including lipid droplets^51^, mitochondria^52^, and protein co-pathologies^53–56^ such as amyloid-b and hyperphosphorylated tau. In *SNCA-*A53T miBrains, the neutral lipid marker Lipid Spot showed significant spatial overlap with both SNCA-A53T-sfGFP and phosphorylated α-Syn. Quantitative volumetric analysis demonstrated a significant increase in Lipid Spot and p-Syn co-localized in *SNCA-*A53T miBrains compared with control miBrains (p = 0.0006; **Figure 2a**), consistent with prior reports describing lipid droplet-associated α-Syn-positive neurotoxic inclusions in human iPSC-derived neurons^29^. The volume of aggregated α-Syn overlapping with mitochondrial marker TOM20 in *SNCA*-A53T miBrains was significantly increased compared to control miBrains (p = 0.017, **Figure S2a**), in agreement with studies demonstrating mitochondrial association of aggregated α-Syn in Lewy pathology^57^. In addition, *SNCA-*A53T miBrains exhibited significantly increased amyloid-b-immunoreactivity (D54D2; p < 0.0001) and hyperphosphorylated tau (PHF1; total: p = 0.0021, **Figure 2b**; somatodendritic: p = 0.0002, **Figure S2b**) compared to controls, suggesting that miBrains can model co-pathologies commonly observed alongside a-Syn inclusion. These data show that α-Syn-rich inclusions in the miBrain are co-associated with lipid, mitochondrial, and proteopathic characteristics of Lewy pathology.

**Figure 2:**
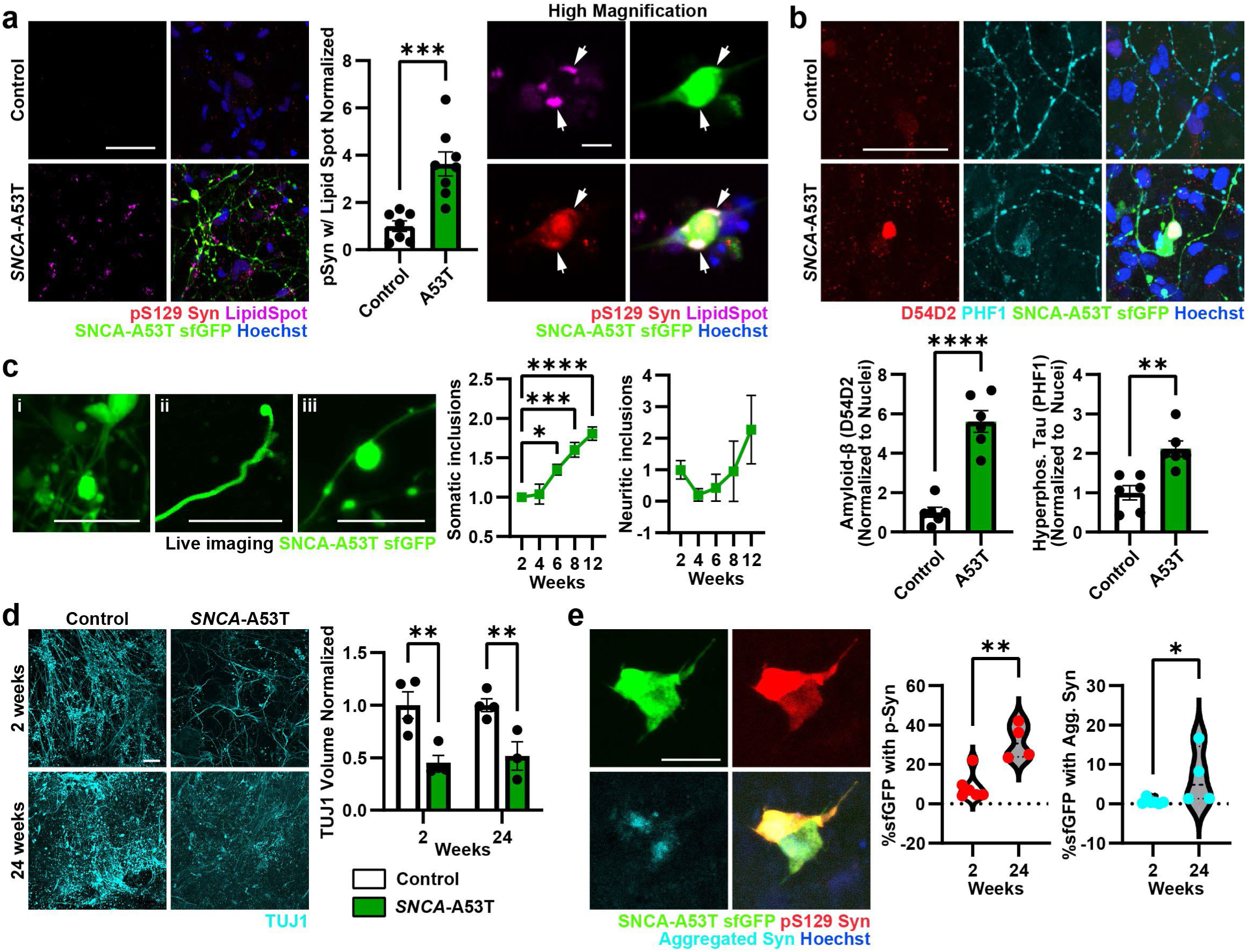
α-Syn inclusions in miBrains recapitulate pathological features of Lewy bodies. **a**. Representative images of SNCA-A53T-sfGFP (green), α-Syn pS129 (red), and LipidSpot (magenta) in control or *SNCA-*A53T 18-day-old miBrains. Arrowhead indicates a lipid droplet within an α-Syn pS129 inclusion. Co-localized α-Syn pS129 and LipidSpot volume was normalized to control. Data are mean ± SEM (n = 8 biological replicates). P-value by two-tailed unpaired t-test. Scale bars: 50 µm (low magnification) and 10 µm (high magnification). **b.** Representative images of amyloid-β (D54D2; red), hyperphosphorylated tau (PHF1; cyan), and SNCA-A53T-sfGFP (green) in control and *SNCA-*A53T 18-day-old miBrains. D54D2 volume and somatic PHF1 were normalized to Hoechst^+^ nuclei (blue) and control. Data are mean ± SEM (n = 6 biological replicates). P-values by two-tailed unpaired t-tests. Scale bar: 50 µm. **c.** Representative live imaging of SNCA-A53T-sfGFP (green) in cell bodies (i), neurites (ii), and varicose-like inclusions (iii). Somatic and neuritic inclusions numbers over time were normalized by sfGFP volume. Data are mean ± SEM (n = 4 biological replicates). P-values by one-way ANOVA and Dunnett’s test. Scale bars: 25 µm (i, ii) and 15µm (iii). **d.** Representative images of control and *SNCA*-A53T miBrains after 2 or 24 weeks in culture. TUJ1 (cyan) volume was normalized to time-matched controls. Data are mean ± SEM (n = 3-4 biological replicates). P-values by 2-way ANOVA and Fisher’s test. Scale bar: 50 µm. **e.** Representative images of *SNCA*-A53T miBrains stained for α-Syn pS129 (red) and aggregated α-Syn after 24 weeks in culture. SNCA-A53T-sfGFP (green) volume co-localized with α-Syn pS129 or aggregated α-Syn was measured in 2- and 24-week miBrains and normalized to 2-week miBrains. Data are mean ± SEM (n = 4-6 biological replicates). P-values by Mann-Whitney test. Scale bar: 25 µm. *p < 0.05, **p < 0.01, ***p < 0.001, ****p < 0.0001. A53T neurons utilized in **c-e** were from the A53T-1A CORR28 line (familial Parkinson’s disease donor).

Lewy bodies are resistant to extraction with non-ionic detergents such as Triton X-100 and become solubilized in stronger detergents such as SDS^58^. To determine whether α-Syn inclusions formed in the miBrain exhibit similar biochemical properties, we performed sequential detergent extraction of the neuronal lysates followed by Western blotting for qualitative assessment of α-Syn species. In control and *SNCA*-A53T 2D neurons, α-Syn was detected exclusively in the Triton-X-100 soluble fraction, with little to no signal in the SDS fraction. In contrast, the *SNCA-*A53T neurons with PFFs showed both total α-Syn and p-Syn in the SDS-soluble fraction in addition to the Triton-soluble pool (**Figure S2c**), demonstrating the presence of detergent-resistant aggregates in the *SNCA*-A53T neurons akin to Lewy Body pathology found in the human brain. These biochemical features are consistent with the detergent solubility profile of Lewy Body-associated proteins^29^. They are also concordant with immunostaining results using p-Syn antibodies in 2D neurons (**Figure S1f**), further supporting the suitability of p-Syn staining as a biomarker for pathogenic a-Syn inclusions.

Lewy pathology comprises dense spherical Lewy Bodies, less dense “pale bodies”, and thread-like or varicose Lewy neurites^44^. Longitudinal live imaging revealed somatic and neuritic SNCA-A53T-sfGFP inclusions in miBrains (**Figure 2c i-iii**), with morphologies resembling human Lewy pathology. Over the first 12 weeks in culture, somatic α-Syn inclusions in miBrains increased significantly (1.81 ± 0.09-fold; p < 0.0001), whereas neuritic inclusions remained stable (p = 0.52; **Figure 2c**), indicating preferential accumulation in neuronal cell bodies over time.

Because α-Syn aggregation is associated with neuronal injury *in vivo*, we next assessed neurotoxicity in miBrains^59,60^. Soluble biomarkers of cell death (lactate dehydrogenase (LDH)) were significantly elevated in the media of *SNCA-*A53T miBrains compared with control (p = 0.026, **Figure S2d**). In parallel, total sfGFP-labeled α-Syn volume significantly declined between weeks 2 and 22 (p < 0.0001, **Figure S2e**), consistent with progressive cell loss. Neuronal content was also reduced in *SNCA-*A53T compared to control miBrains, as measured by TUJ1-positive volume at both 2 weeks (p = 0.002) and 24 weeks (p = 0.007; **Figure 2d**). Over this interval, the fraction of sfGFP signal co-localizing with pS129 α-Syn (p = 0.0095) and aggregated α-Syn (p = 0.0381) increased (**Figure 2e**), consistent with increased aggregation of α-Syn over time. Collectively, these results demonstrate that miBrains phenocopy morphologically and biochemically complex α-Syn pathology accompanied by neuronal injury.

### APOE4 increases α-Syn phosphorylation and aggregation through non-cell autonomous mechanisms in miBrain tissue

*APOE4* is a genetic risk factor for α-Syn in LBD^14^ and AD^61,62^, and is associated with increased disease severity^63,64,65^. To investigate how *APOE4* promotes α-Syn pathology, we used isogenic iPSC lines derived from an *APOE3/3* donor and CRISPR-edited to *APOE4/4*^66,67^. In monocultures, *APOE4/4 SNCA-*A53T neurons exhibited comparable levels of p-Syn to isogenic *APOE3/3 SNCA*-A53T neurons, both with and without PFF exposure (**Figure S3a**), indicating no detectable neuronal cell-autonomous effects of *APOE* genotype under these conditions. We therefore hypothesized that *APOE4* enhances α-Syn pathology through non-neuronal cell types. Using the same isogenic lines, we generated astrocytes, OPCs, endothelial cells, mural cells, and microglia, confirming comparable cell type-specific markers across *APOE* genotypes for each cell type (**Figure S3b**). We then assembled isogenic *APOE3/3* and *APOE4/4* miBrains containing *APOE3/3 SNCA*-A53T or *APOE4/4 SNCA*-A53T neurons respectively. Consistent with clinical studies, *APOE4/4* miBrains displayed significantly increased neuronal p-Syn compared to *APOE3/3* miBrains (p = 0.0001, **Figure 3a**). The volume of p-Syn outside the sfGFP signal was also significantly increased in *APOE4/4* miBrains (p = 0.0263, **Figure S3c**), indicating enhanced phosphorylation of endogenous (non-A53T) α-Syn in the *APOE4/4* miBrains compared to isogenic controls. To assess whether increased phosphorylation of α-Syn in the *APOE4/4* miBrain is a direct cell-autonomous effect of *APOE4/4* neurons, we generated miBrains that were all *APOE4/4* except for *APOE3/3 SNCA*-A53T neurons. Strikingly, we observed a significant increase in p-Syn even when *APOE3/3 SNCA*-A53T neurons were placed in an otherwise *APOE4/4* miBrain (p < 0.0001, **Figure S3d**). These results demonstrate that *APOE4/4* promotes neuronal α-Syn pathological phenotypes through non-cell autonomous mechanisms within a multicellular human brain-like tissue environment.

**Figure 3.**
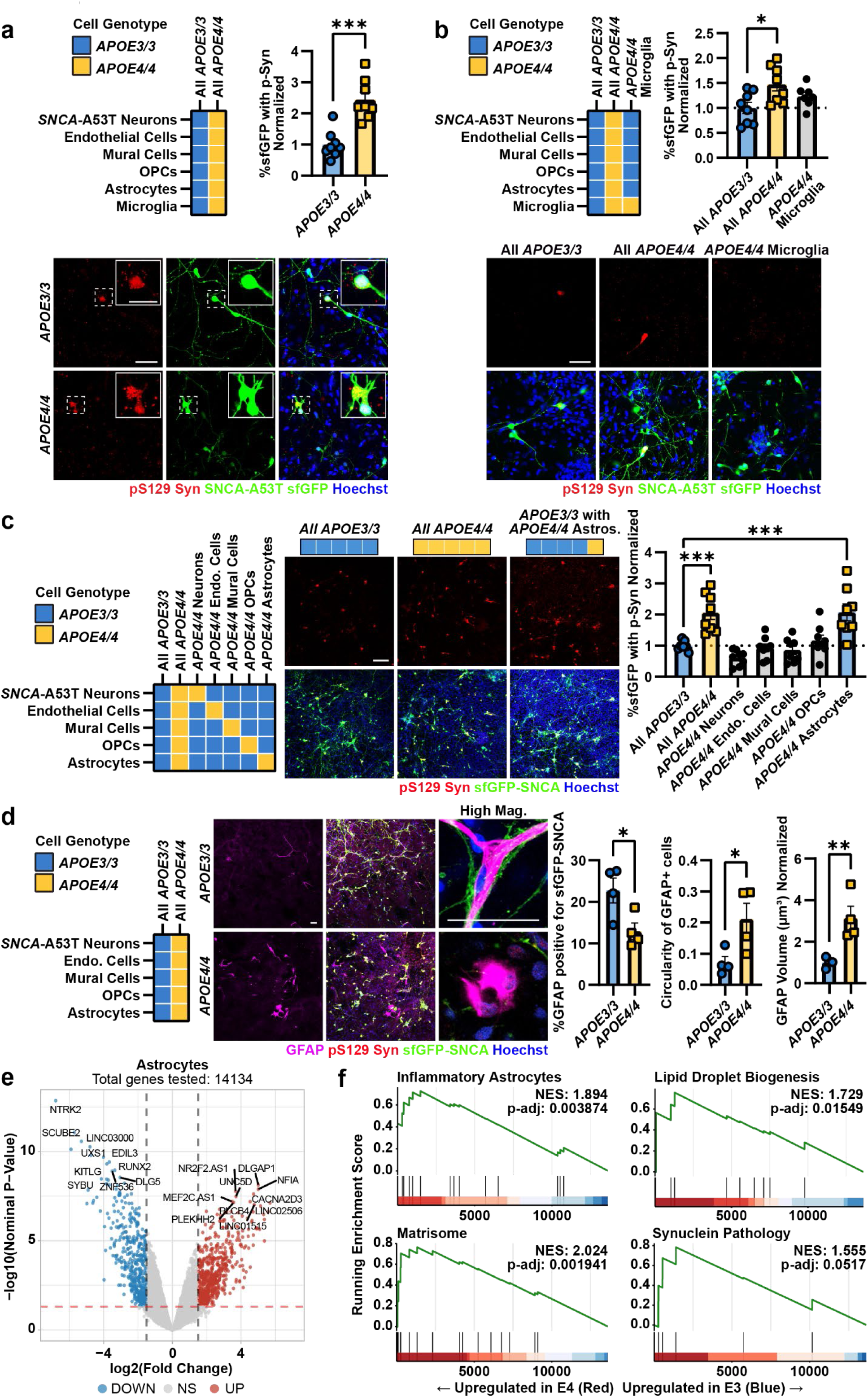
*APOE4* promotes α-Syn phosphorylation through astrocytes. **a.** Representative images of *APOE3/3* and *APOE4/4* 18-day-old miBrains with SNCA-A53T-sfGFP (green) neurons stained for α-Syn pS129 (red). α-Syn pS129 volume within SNCA-A53T-sfGFP neurons was normalized to total SNCA-A53T-sfGFP volume and *APOE3/3*. Data are mean ± SEM (n = 8 biological replicates). P-value by two-tailed unpaired t-test. Scale bar: 50 µm (low magnification) and 25 µm (insets). **b.** Representative images of *APOE3/3, APOE4/4,* and *APOE3/3* with *APOE4/4* microglia 18-day-old miBrains stained for α-Syn pS129 (red). α-Syn pS129 volume within SNCA-A53T-sfGFP neurons (green) was normalized to total SNCA-A53T-sfGFP volume and *APOE3/3*. Data are mean ± SEM (n = 8 biological replicates). P-values by one-way ANOVA and Tukey’s test. Scale bar: 50 µm. **c.** Representative images of *APOE3/3, APOE4/4,* and *APOE3/3* with *APOE4/4* astrocytes 18-day-old miBrains (without microglia) stained for α-Syn pS129 (red). α-Syn pS129 volume within SNCA-A53T-sfGFP neurons (green) was normalized to total SNCA-A53T-sfGFP volume and *APOE3/3*. Data are mean ± SEM (n = 8 biological replicates). P-values by one-way ANOVA and Dunnett’s test. Scale bar: 50 µm. **d.** Representative images of *APOE3/3* and *APOE4/4* 18-day-old miBrains (without microglia) stained for of GFAP (magenta). Quantification of GFAP overlap with SNCA-A53T-sfGFP (green; left), circularity (center), and GFAP volume normalized to *APOE3/3* (right). Data are mean ± SEM (n = 4 biological replicates). P-values by two-tailed unpaired t-tests. Scale bar: 50 µm. **e.** Volcano plot of differentially expressed genes (DEGs) in *APOE4/4* vs. *APOE3/3* astrocytes (upregulated, red; downregulated, blue; |log2 fold change| > 1.5, p-value < 0.05). **f.** Gene set enrichment analysis (GSEA) of *APOE4/4* versus *APOE3/3* astrocytes. *p < 0.05, **p < 0.01, ***p < 0.001, ****p < 0.0001

### APOE4/4 astrocytes mediate increased α-Syn pathological phenotypes in miBrains

Our results suggest that non-neuronal cell types are responsible for increased p-Syn in the *APOE4/4* miBrains. Because microglia are implicated in α-Syn clearance^68,69^ and have been reported to enhance tau hyperphosphorylation in *APOE4* brain tissue^27^, we first examined whether microglia modulate α-Syn pathology in our system. In 2D monocultures, microglia exposed to conditioned media from *SNCA-*A53T neurons showed increased expression of activation and phagocytic markers, including IBA1 (p = 0.0553) and CD68 (p = 0.0122, **Figure S3e**), consistent with microglial activation and phagocytic activity. However, inclusion of microglia in miBrains did not significantly alter p-Syn levels compared to miBrains without microglia for either *APOE3/3* (p = 0.943) or *APOE4/4* (p = 0.585) miBrains (**Figure S3f**). To further test microglial genotype effects, we replaced *APOE3/3* microglia with *APOE4/4* microglia in an otherwise *APOE3/3* miBrain and observed no significant increase in α-Syn phosphorylation (p = 0.349; **Figure 3b)**. Together, these results demonstrate that microglia, regardless of their *APOE* genotype, are not a major determinant of α-Syn phosphorylation in miBrains.

To identify which *APOE4/4* cell types promote increased α-Syn pathology, we generated a series of isogenic hybrid miBrains in which individual *APOE3/3* cell types were selectively substituted with isogenic *APOE4/4* counterparts (**Figure 3c**). Based on the results above (**Figure 3b and S3f**), microglia were excluded to reduce combinatorial complexity. Consistent with previous results (**Figure 3a**), all *APOE4/4* miBrains had significantly increased p-Syn staining compared to isogenic all *APOE3/3* miBrains (p = 0.0005). Replacement of *APOE3/3* neurons, endothelial cells, mural cells, or OPCs with *APOE4/4* isogenic cells did not significantly increase p-Syn levels in the miBrain (**Figure 3c**). In contrast, selective substitution of *APOE4/4* astrocytes into an otherwise *APOE3/3* miBrain was sufficient to significantly increase in p-Syn immunoreactivity (p = 0.0004), reaching levels comparable to *APOE4/4* miBrains (**Figure 3c**). Collectively these results demonstrate that *APOE4/4* astrocytes are responsible for increasing α-Syn pathology in *APOE4/4* miBrain.

Because astrocytes contribute to neuronal α-Syn uptake and degradation, we next examined astrocyte-α-Syn interactions^70–72^. Compared to isogenic *APOE3/3* miBrains, in *APOE4/4 miBrains* GFAP-positive astrocytes showed significantly reduced spatial overlap with SNCA-A53T-sfGFP (p = 0.0348) and displayed a more amoeboid reactive morphology characterized by increased circularity (p = 0.0398) and elevated GFAP expression (p = 0.010 (**Figure 3d and Videos S2 and S3**). snRNA-seq analysis of *APOE3/3* and *APOE4/4* miBrains showed that *APOE4/4* astrocytes exhibit transcriptional alterations consistent with altered astrocytic α-Syn handling associated with increased reactivity (**Figure 3e**). In *APOE4*/4 astrocytes the number of DEGs (1007 DEGs, adjusted p< 0.05, |logFC| > 0.5) was more than double that of *APOE4/4* neurons harboring *SNCA*-A53G (420 DEGs) when compared to their *APOE3/3* counterparts (**Table S1**). GSEA of astrocytes showed altered expression of genes associated with inflammatory states (P = 0.03), matrisome (P = 0.0004), α-Syn pathology (P = 0.04), and lipid droplets biogenesis (P = 0.04) in *APOE4/4* astrocytes (**Figure 3f, Table S2**). These findings indicate that the *APOE4/4* genotype promotes astrocyte activation consistent with astrogliosis and enhanced responses to α-Syn pathology and/or the miBrain environment. Collectively, these results demonstrate that astrocytes play a dominant non-cell-autonomous role in promoting neuronal α-Syn phosphorylation and aggregation in miBrains.

### APOE4/4 astrocytes exhibit impaired uptake and degradation of exogenous α-Syn

To define the mechanisms by which *APOE4/4* astrocytes increase the phosphorylation and aggregation of neuronal α-Syn, we first asked whether increased astrocytic *SNCA* expression contributes to the increased α-Syn burden in the *APOE4/4* miBrain and human brain tissue. Analysis of single-nucleus transcriptomics data from post-mortem human brain^73^ showed that *SNCA* mRNA is significantly reduced in astrocytes from *APOE4* carriers (*APOE3/4* and *APOE4/4*) compared to control age-matched *APOE3/3* individuals (p = 0.019; **Figure 4a**). We observed a similar decrease in *SNCA* transcript levels in isogenic *APOE4/4* versus *APOE3/3* iPSC-derived astrocytes^66^ (p = 0.017) (**Figure S4a**), indicating that transcriptional upregulation of *SNCA* in astrocytes is unlikely to account for increased α-Syn pathology in *APOE4* contexts. In contrast, immunoblotting of astrocyte monocultures revealed increased total and phosphorylated α-Syn protein in *APOE4/4* astrocytes compared with isogenic *APOE3/3* astrocytes across two donors (Total α-Syn: Donor #1: p = 0.023; Donor #2: p= 0.043 and phosphorylated α-Syn; Donor #1: p = 0.0047; Donor #2: p = 0.0137; **Figure 4b; Figure S4b**) suggesting that post-translational mechanisms underlie increased α-Syn abundance in *APOE4/4* astrocytes.

**Figure 4.**
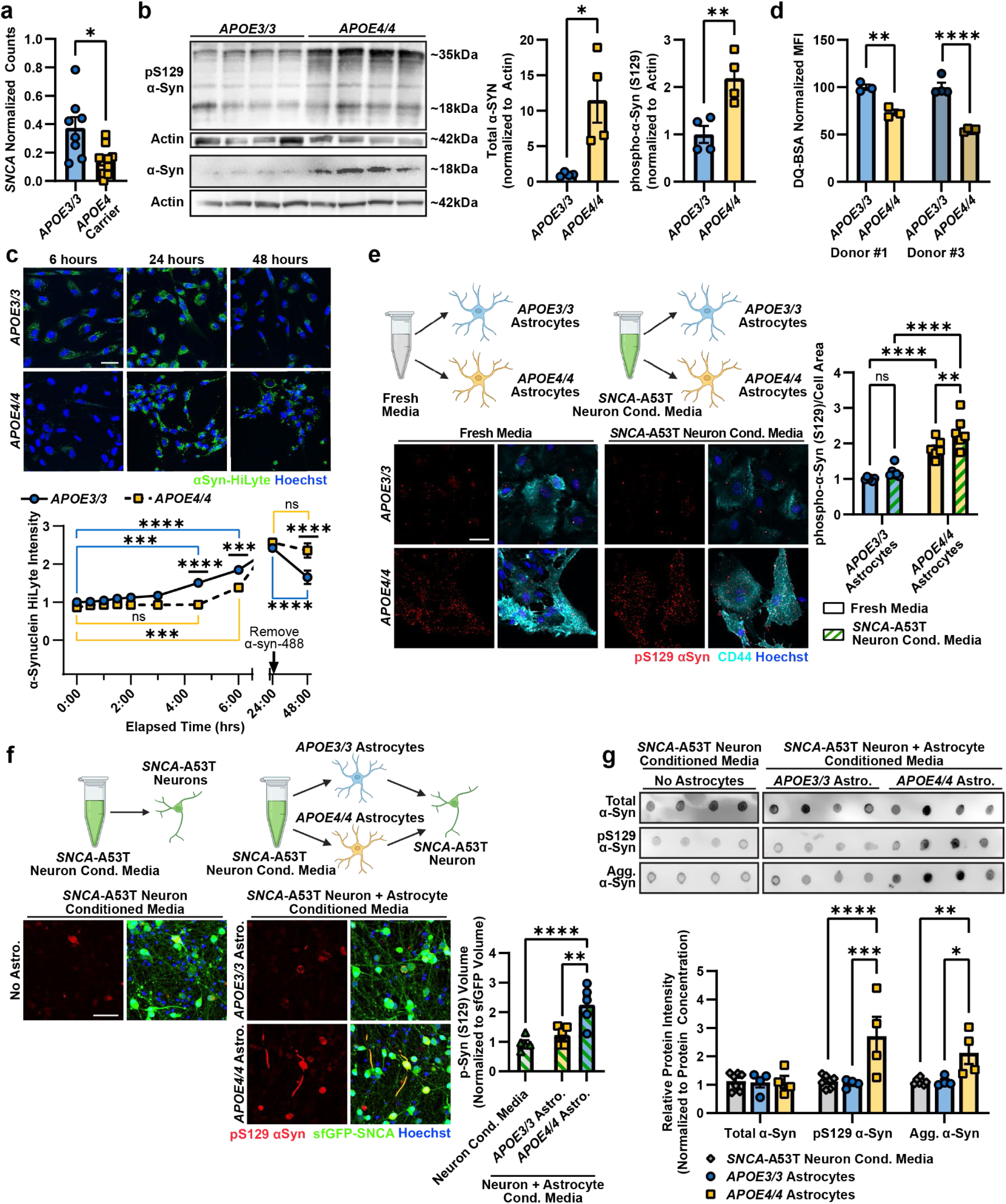
Impaired lysosomal function in *APOE4/4* astrocytes seeds neuronal α-Syn phosphorylation. **a.** Pseudobulk analysis of *SNCA* expression in human astrocytes from Haney et al. 2024, normalized to gene count (TPM). Bars represent mean ± SEM (n = 8 *APOE3/3*, n = 10 *APOE4* carriers). P-value by two-tailed unpaired t-test. **b** Western blots of *APOE3/3* and *APOE4/4* astrocytes for total and phosphorylated α-Syn, and β-Actin. Quantification was normalized to β-Actin and *APOE3/3.* Data are mean ± SEM (n = 4 replicates). P-values by Welch’s one-tailed t-test **c.** Representative images of uptake and degradation of fluorescent α-Syn-HiLyte (green). α-Syn-HiLyte was removed after 24 hours, and fluorescence intensity was normalized to Hoechst (blue) area and *APOE3/3* at the first time point. Data are mean ± SEM (n = 4 biological replicates). P-values by 2-way ANOVA and Tukey’s test. Scale bar = 50 µm. **d.** DQ-BSA Green mean fluorescence intensity measured in DAPI negative cell population by flow cytometry after 24 hours in two isogenic iPSC lines. Data are mean ± SEM (n = 3-4 biological replicates). P-values by two-tailed, unpaired t-tests. **e.** Representative images of pS129 α-Syn (red) in *APOE3/3* and *APOE4/4* astrocytes after exposure to fresh or conditioned neuron media. pS129 α-Syn mean intensity was normalized to cell area (CD44; cyan) and *APOE3/3* astrocytes exposed to fresh media. Data are mean ± SEM (n = 6 biological replicates). P-values by 2-way ANOVA and Fisher’s LSD. Scale bar = 50 µm. **f.** Representative images of pS129 α-Syn (red) in *SNCA*-A53T neurons treated with neuron-conditioned media (left) or neuron and astrocyte-conditioned media from either *APOE3/3* or *APOE4/4* astrocytes (right). pS129 α-Syn volume was normalized to *SNCA-*A53T-sfGFP (green) volume and neuron-conditioned media samples. Data are mean ± SEM (n = 5 biological samples). P-values by one-way ANOVA and Tukey’s test. Scale bar = 50 µm. **g.** Dot blots of neuron-conditioned media or double-conditioned media from SNCA-A53T neurons and isogenic *APOE3/3* or *APOE4/4* astrocytes for total, phosphorylated, and aggregated αSyn. All dots were on one blot; the image was cropped for presentation. Signal was normalized to protein concentration measured by BCA assay. Data are mean ± SEM (n = 4 biological replicates). P-values by 2-way ANOVA and Tukey’s test. *p < 0.05, **p < 0.01, ***p < 0.001, ****p < 0.0001

Astrocytes have a well-described protective and homeostatic function of taking up and degrading neuronal α-Syn^70–72^. Given this role and our findings that *APOE4/4* astrocytes increase neuronal α-Syn phosphorylation and co-localize less with SNCA-A53T-sfGFP in the miBrain (**Figure 3d**), we tested whether *APOE4/4* astrocytes have impaired processing of exogenous α-Syn. Isogenic *APOE3/3* and *APOE4/4* astrocytes were exposed to fluorescently labeled α-Syn-HiLyte, and uptake and degradation were monitored by live imaging. *APOE3/3* astrocytes showed rapid uptake of α-Syn with significant intracellular accumulation by 4.5 hours (p = 0.001) and 6 hours (p < 0.0001) (**Figure 4c**). In contrast, isogenic *APOE4/4* astrocytes showed delayed uptake, with no detectable increase of α-Syn at 4.5 hours (p = 0.9997) and only a modest increase by 6 hours (p = 0.0006) (**Figure 4c**). To assess degradation, extracellular α-Syn was removed after 24 hours, and intracellular α-Syn-HiLyte signal was tracked for an additional 24 hours. *APOE3/3* astrocytes showed a significant reduction in intracellular α-Syn-HiLyte signal (p < 0.0001), whereas *APOE4/4* astrocytes showed no significant decrease (p = 0.694; **Figure 4c**), indicating impaired degradation of α-Syn. Together, these results demonstrate that *APOE4/4* astrocytes accumulate α-Syn protein despite lower *SNCA* transcript levels and exhibit defects in both uptake kinetics and intracellular clearance of exogenous α-Syn.

### APOE4/4 astrocytes exhibit impaired lysosomal proteolytic function

Because intracellular protein turnover is primarily mediated by lysosomal and proteasomal pathways^74^, we next investigated which pathway is primarily responsible for α-Syn degradation in astrocytes. We treated the cells with the lysosomal inhibitor bafilomycinA1 or the proteasomal inhibitor MG-132 and monitored the degradation of fluorescent α-Syn-HiLyte. Lysosomal inhibition significantly blocked the degradation of α-Syn-HiLyte (p = 0.0241), whereas proteasomal inhibition had no significant effect (p = 0.079; **Figure S4c**), consistent with prior reports that astrocytic α-Syn clearance occurs predominantly through the endolysosomal pathway^75^. These findings support a model in which impaired lysosomal function promotes α-Syn accumulation.

We therefore tested whether *APOE4/4* astrocytes exhibit endolysosomal dysfunction relative to isogenic *APOE3/3* astrocytes. Lysosomal proteolytic capacity was assessed using DQ-BSA, a self-quenched substrate that fluoresces upon lysosomal cleavage. *APOE4/4* astrocytes showed significantly reduced lysosomal DQ-BSA signal compared to isogenic control *APOE3/3* astrocytes across two isogenic donor pairs (Donor #1: p= 0.0032; Donor #2: p < 0.0001, **Figure 4d**), indicating decreased lysosomal proteolytic activity. Consistent with this, immunoreactivity for the lysosomal membrane protein LAMP1 was also significantly reduced in *APOE4/4* astrocytes compared to isogenic *APOE3/3* astrocytes (Donor #1: p = 0.0002 and Donor #2: p = 0.0001, **Figure S4d**). To distinguish between reduced lysosomal number and reduced lysosomal function per organelle, we normalized DQ-BSA fluorescence to co-internalized dextran, which accumulates in the lysosomes but is resistant to proteolysis^76^. Even after dextran normalization, *APOE4/4* astrocytes retained significantly lower proteolytic activity (p = 0.0098; **Figure S4e**), indicating functional impairment rather than solely reduced lysosome abundance. As expected, bafilomycinA1 suppressed DQ-BSA cleavage in both *APOE3/3* (p < 0.0001) and *APOE4/4* astrocytes (p = 0.0174; **Figure S4e**), confirming lysosome-dependent signal generation. Collectively, these results demonstrate that *APOE4/4* astrocytes exhibit both reduced lysosomal mass and diminished lysosomal proteolytic function, providing a mechanistic basis for impaired lysosomal clearance of a-Syn.

### APOE4 astrocytes take up neuronal α-synuclein and release modified pathogenic seeds

Since phosphorylated α-Syn is a key readout of α-Syn pathology, we next determined whether astrocytes can phosphorylate neuronal α-Syn. Therefore, we exposed Isogenic *APOE3/3* and *APOE4/4* astrocytes to fresh media or conditioned media from *SNCA-*A53T neurons. At baseline, *APOE4/4* astrocytes exhibited higher endogenous p-Syn levels than *APOE3/3* astrocytes (**Figure 4e**; p < 0.0001), consistent with our immunoblotting results (**Figure 4b**). *APOE4/4* astrocytes incubated with media previously exposed to SNCA-A53T neurons for 3 days showed a further increase in p-Syn (p = 0.007) while APOE3/3 astrocytes were unaffected (p = 0.232) (**Figure 4e**), indicating that *APOE4/4* astrocytes increase α-Syn phosphorylation. However, in the miBrain and post-mortem human brain, inclusions of α-Syn are primarily found inside neurons. We found that cell culture media collected from *SNCA-*A53T neurons and then inoculated onto fresh *SNCA-*A53T neuronal cultures itself does not increase phosphorylation of neuronal α-Syn (**Figure 4f**), suggesting that non-cell-autonomous mechanisms likely modify α-synuclein’s pathogenicity. Therefore, we hypothesized that *APOE4/4* astrocytes fail to degrade α-Syn due to impaired endolysosomal function. Instead, *APOE4/4* astrocytes phosphorylate α-Syn and release the more pathogenic forms of a-Syn into the extracellular space allowing them to be taken up by neurons and promote the formation of neuronal α-Syn inclusions. To test this hypothesis, we first exposed *SNCA-*A53T neurons to conditioned media from only *APOE3/3* or *APOE4/4* astrocytes. Astrocyte-conditioned media itself did not induce significant p-Syn in *SNCA-*A53T neurons, and no significant difference was observed between *APOE3/3* and *APOE4/4* conditioned media (*APOE3/3*: p = 0.9903; *APOE4/4*: p = 0.9092; **Figure S4f**). We reasoned the low levels of α-Syn produced by astrocytes alone is not sufficient to induce phosphorylation and aggregation of neuronal α-Syn. Therefore, we performed a double conditioned media experiment where conditioned media was first collected from *SNCA-*A53T neurons and subsequently cultured with either *APOE3/3* or *APOE4/4* astrocytes. This double-conditioned media was then collected and inoculated onto fresh *SNCA-*A53T neuron monocultures (**Figure 4f**). Neurons grown in *APOE3/3* double-conditioned media did not show a significant increase in α-Syn phosphorylation (p = 0.601). In contrast, *APOE4/4* double-conditioned media led to a 2.25-fold increase (± 0.30 SEM) in α-Syn phosphorylation (p < 0.0001, **Figure 4f**). To assess whether the induction of p-Syn by *APOE4/4* double-conditioned media was a failure of astrocytes to degrade α-Syn or a modification of α-Syn, we analyzed the α-Syn species in the double-conditioned media via dot blot. Although the total levels of α-Syn in the media were not different between *APOE3/3* and *APOE4/4* astrocytes (p = 0.999), p-Syn and aggregated α-Syn were both significantly increased (p = 0.0002 and p = 0.0232) in media conditioned by *APOE4/4* astrocytes compared to *APOE3/3* astrocytes (**Figure 4g**). *APOE3/3* astrocyte double-conditioned media appeared not to change having similar p-Syn and aggregated α-Syn levels to the original *SNCA*-A53T media that was not conditioned by astrocytes (p = 0.971 and p = 0.999; **Figure 4g**).

To determine whether these pathogenic α-Syn species originated from astrocytic endogenous α-Syn or from neuronal-derived α-Syn processed by astrocytes, we generated *APOE4/4* astrocytes with >90% *SNCA* knockdown using two independent shRNA constructs (shRNA#1: 94.25% ± 0.55 SEM knockdown efficiency; shRNA#2: 94.70% ± 1.5 SEM knockdown efficiency; **Figure S4g**). *SNCA* knockdown in *APOE4/4* astrocytes did not reduce the ability of double-conditioned media to induce neuronal p-Syn compared to control *APOE4/4* astrocytes (no shRNA and scrambled shRNA; shRNA #1: p = 0.997, shRNA #2: p = 0.998; **Figure S4h**). Collectively, these results indicate that *APOE4/4* astrocytes do not contribute to pathology through the secretion of their endogenous α-Syn. Instead, *APOE4/4* astrocytes take up neuronal-derived α-Syn, promote its pathogenic modification, and release seeding-competent α-Syn that promotes neuronal α-Syn inclusion formation.

### Cholesterol accumulation in APOE4/4 astrocytes leads to lysosomal dysfunction and pathogenic α-Syn processing

We next investigated how APOE4 in astrocytes leads to lysosomal dysfunction and the spread of pathogenic α-Syn. In the brain, APOE is primarily expressed in astrocytes and regulates cholesterol transport to neurons to support membrane and synapse homeostasis^77–79^. APOE4 is associated with reduced lipid transport capacity^80^, cholesterol accumulation within astrocytic lysosomes, and impaired lysosomal function^67,81^. In agreement with previous reports, transcriptomic analysis of *APOE4/4* astrocytes showed differential expression of genes (p < 0.05) involved in lipid storage and transport, as well as response to lipopolysaccharide and oxidative stress, which are associated with a neurotoxic reactive phenotype (**Figure S5a**)^67,82,83^. Consistent with altered lipid metabolism, *APOE4/4* astrocytes showed significantly increased BODIPY-cholesterol accumulation compared to isogenic *APOE3/3* astrocytes generated from two different individuals (Donor #1: p = 0.001; Donor #2: p = 0.002, **Figure 5a**). Analysis of *APOE4/4* astrocytes within miBrains by snRNA-seq indicate an enrichment of gene programs associated with lipid droplet biogenesis (**Figure 3f, Table S2**).

**Figure 5.**
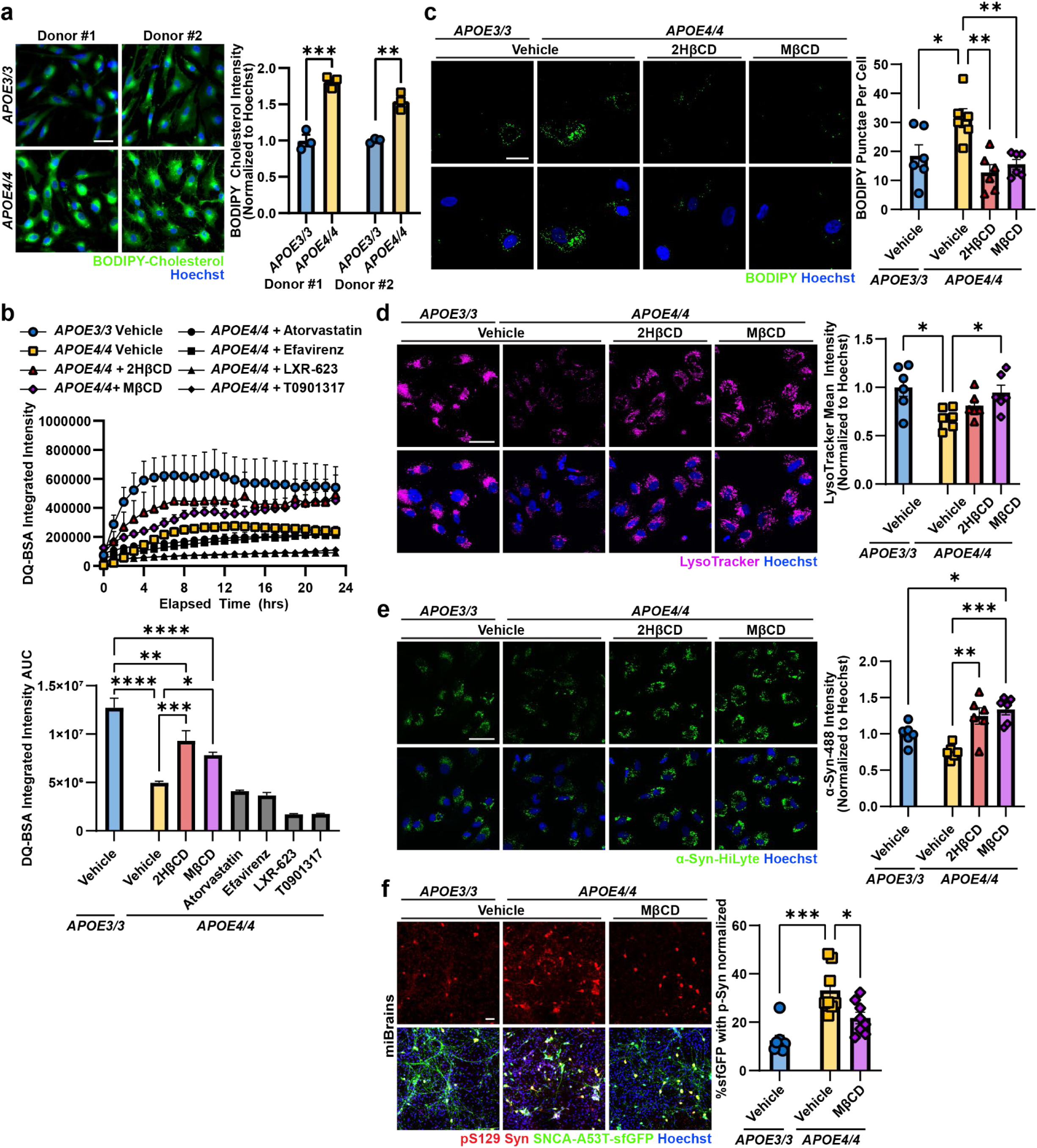
Cholesterol reduction restores lysosomal function and α-Syn homeostasis in *APOE4/4* astrocytes. **a.** Representative images of live astrocytes with BODIPY-cholesterol (green) in two isogenic lines. BODIPY-cholesterol mean intensity was normalized to Hoechst (blue) area and *APOE3/3*. Data are mean ± SEM (n= 3 biological replicates). P-values by two-tailed unpaired t-tests. Scale bar = 50 µm. **b.** Top: DQ-BSA Red integrated intensity over 24 hours in astrocytes treated with 2HβCD, MβCD, atorvastatin, efavirenz, LXR-623, or T0901317. Data are mean ± SEM (n = 4 biological replicates). Bottom: area under the curve. Data are mean ± SEM (n = 4 biological replicates). P-values by 2-way ANOVA and Tukey’s test. **c.** Representative images of BODIPY 493/503 (green) staining in astrocytes treated with 2HβCD or MβCD. BODIPY puncta were normalized to Hoechst (blue) count. Data are mean ± SEM (n = 6 biological replicates). P-values by 2-way ANOVA and Tukey’s test. **d.** Representative images of LysoTracker (magenta) in astrocytes treated with 2HβCD or MβCD. LysoTracker mean intensity was normalized by Hoechst (blue) area and *APOE3/3*. Data are mean ± SEM (n = 5-6 biological replicates). P-values by 2-way ANOVA and Sidak test. Scale bar = 50 µm. **e.** Representative images of astrocytes treated with cyclodextrins after a 24h incubation with fluorescent α-Syn-HiLyte (green), in two isogenic lines. α-Syn-HiLyte intensity was normalized to Hoechst (blue) area and *APOE3/3*. Data are mean ± SEM (n = 6 biological replicates). P-values by 2-way ANOVA and Tukey’s test. Scale bar = 50 µm. **f.** Representative images of *APOE3/3*, *APOE4/4*, and *APOE4/4* treated with MβCD miBrains stained for pS129 α-Syn (red). pS129 α-Syn volume within SNCA-A53T-sfGFP (green) normalized to total sfGFP volume and *APOE3/3* 18-day-old miBrains. Data are mean ± SEM (n = 8 biological replicates). P-values were calculated by 2-way ANOVA and Tukey’s test. Scale bar = 50 µm. *p < 0.05, **p < 0.01, ***p < 0.001, ****p < 0.0001.

Given the coexistence of cholesterol accumulation and lysosomal impairment, we hypothesized that excess intracellular cholesterol contributes to impaired α-Syn degradation. We conducted a primary screen for improved lysosomal proteolytic activity, as measured by DQ-BSA cleavage, in *APOE4/4* astrocytes treated with small molecules known to reduce cholesterol bioavailability. Treatment with 2-hydroxypropyl-β-cyclodextrin (2HβCD) and methyl-β-cyclodextrin (MβCD) significantly increased lysosomal proteolysis (2HβCD: p = 0.0002; MβCD: p = 0.019) (**Figure 5b**), whereas the other interventions had neutral or negative effects on *APOE4/4* lysosomal proteolytic activity (**Figure 5b**). Treatment with cyclodextrins significantly decreased neutral lipids in *APOE4/4* astrocytes as measured by BODIPY staining (2HβCD: p = 0.0013; MβCD: p = 0.0063) to levels comparable to *APOE3/3* astrocytes (**Figure 5c**). Filipin staining revealed a significant increase in endogenous free cholesterol in *APOE4/4* astrocytes compared to *APOE3/3* astrocytes, which was significantly reduced following MβCD treatment (p = 0.0167) to levels comparable to *APOE3/3* astrocytes (**Figure S5b**). Increased lysosomal proteolytic activity with MβCD was replicated in astrocytes from a second donor (p = 0.0045; **Figure S5c**).

Consistent with improved lysosomal capacity, MβCD increased LysoTracker-positive lysosomal signal in *APOE4/4* astrocytes (p = 0.044; **Figure 5d**), with similar effects observed across donors (2HβCD: p < 0.0001; MβCD: p < 0.0001, **Figure S5d**). We next tested whether cholesterol reduction improves astrocytic α-Syn processing. Cyclodextrin-treated *APOE4/4* astrocytes showed significantly increased uptake of α-Syn-HiLyte at 24 hours (2HβCD: p = 0.0009; MβCD: p = 0.0001) compared to vehicle-treated *APOE4/4* astrocytes (**Figure 5e**). This effect was replicated in isogenic astrocytes from a second individual (2HβCD: p = 0.0002; MβCD: p = 0.0001) (**Figure S5e**). To validate that the α-Syn was being endocytosed into the endolysosomal pathway, we incubated astrocytes with α-Syn-HiLyte for 4 hours and co-stained with LysoTracker. In agreement with our other findings that *APOE4/4* astrocytes have impaired α-Syn uptake, the percentage of α-Syn-HiLyte that co-localized with LysoTracker decreased in *APOE4/4* astrocytes compared to *APOE3/3* (p = 0.0395; **Figure S5f**). However, *APOE4/4* astrocytes treated with MβCD showed increased colocalization of α-Syn-HiLyte with LysoTracker compared to vehicle-treated *APOE4/4* astrocytes (p = 0.0343). Strikingly *APOE4/4* astrocytes treated with MβCD showed similar α-Syn lysosomal localization as vehicle-treated *APOE3/*3 astrocytes (p = 0.946; **Figure S5f**). Collectively, these results indicate that intracellular cholesterol accumulation impairs lysosomal uptake and degradation of α-Syn in *APOE4/4* astrocytes and that extracting cholesterol with cyclodextrin restores these functions.

Our findings implicate astrocytes as a critical cell type in α-Syn processing in the genetic context of *APOE4.* However, α-Syn pathology in Lewy body diseases is characterized by inclusions found within neuronal projections and soma. To investigate whether improved lysosomal activity in cyclodextrin-treated astrocytes affected neuronal α-Syn phosphorylation, we returned to the miBrain model. After 7 days of treatment with MβCD, the levels of neuronal phosphorylated α-Syn were significantly reduced in MβCD-treated *APOE4/4* miBrains (p = 0.0172) when compared to untreated *APOE4/4* miBrains and restored phosphorylation to levels that were not statistically different from *APOE3/3* miBrains (p = 0.059) (**Figure 5f**). These results show that MβCD treatment reduces α-Syn phosphorylation in neurons, likely via restoration of lysosomal activity and α-Syn processing in astrocytes, alleviating cells of the cytotoxic burden of α-Syn aggregation. This may open a therapeutic avenue for cholesterol-lowering pharmacological interventions in the treatment of synucleinopathies and AD with Lewy Bodies, particularly in *APOE4* carriers.

## Discussion

By combining stem cell and genetic engineering, we extended the miBrain platform^27,84^ to model α-Syn pathology and dissect disease-relevant mechanisms. Multicellular miBrains developed α-Syn pathological phenotypes, cell-type-specific transcriptional responses, and neuronal loss. Using genetically hybrid miBrains, we identified astrocytes as the principal mediators of *APOE4*-dependent neuronal α-Syn pathology. Our results establish a causal link between cholesterol dysregulation in *APOE4/4* astrocytes and α-Syn pathology with translational implications for synucleinopathies and AD with Lewy Bodies.

Cholesterol homeostasis is tightly regulated under physiological conditions^85–88^. *APOE4/4* astrocytes exhibit impaired cholesterol efflux^67^ and lysosomal cholesterol acculmulation^66,81,89,90^, consistent with our observation of increased free cholesterol and neutral lipids, and with recent lipidomic analyses of the human brain reporting increased cholesteryl esters in *APOE4* astrocytes^91^. Cholesterol accumulation has been associated with endolysosomal dysfunction^81^, and we speculate that, in *APOE4/4* astrocytes, impaired cholesterol trafficking from late endosomes/lysosomes to downstream compartments promotes sequestration of free cholesterol within the endolysosomal system, secondary accumulation of esterified cholesterol and other neutral lipids into lipid droplets, and reduced lysosomal proteolysis. Consistent with this model, *APOE4* astrocytes displayed reduced lysosomal proteolytic capacity and decreased LAMP1 signal, whereas cyclodextrins reduced intracellular lipid burden, restored lysosomal function, and rescued α-Syn processing. Collectively, these findings support cholesterol trafficking defects as an upstream mediator of astrocytic lysosomal dysfunction.

Astrocytes normally contribute to extracellular α-Syn clearance through uptake and degradation of neuron-derived protein. We show that *APOE4/4* astrocytes instead shift towards pathological α-Syn handling, incompletely degrading internalized α-Syn, promoting its pathogenic modification, and releasing species with enhanced neuron seeding. Although neuronal secretion and astrocytic uptake of α-Syn have been described ^70–72,92^, the conversion and release of pathogenic α-Syn species by dysfunctional astrocytes has not, to our knowledge, been directly demonstrated. Cholesterol accumulation has been linked to increased extracellular vesicle release^93^, raising the possibility that lipid-loaded *APOE4/4* astrocytes adopt a secretory inflammatory state that may contribute to α-Syn propagation. This is consistent with the inflammatory and morphogenic transcriptional signatures we identified in *APOE4/4* astrocytes by snRNA-seq. Additionally, lysosomal lipid imbalance is a convergent feature across synucleinopathies^94–98^, and cholesterol accumulation may directly promote α-Syn aggregation^99,100^ upstream of phosphorylation^101–105^, which itself is not exclusively pathological^106^. Defining the carriers and conformations of astrocyte-released α-Syn species will be an important direction for future work.

Therapeutically, we show that pharmacological reduction of intracellular cholesterol with methyl-b-cyclodextrin reduces neuronal α-Syn phosphorylation in *APOE4/4* miBrains, linking correction of astrocytic lipid imbalance to downstream neuronal benefit. Cyclodextrins are currently under clinical investigation for Neimann-Pick disease Type C, where early studies suggest symptomatic benefits with manageable toxicity^107^. More broadly, cholesterol-lowering strategies have shown mixed results in synucleinopathy models and clinical studies^108,109^. Our results suggest that *APOE* genotype and cholesterol handling may influence therapeutic responsiveness, raising the possibility that genetically stratified lipid-targeting therapies could be more effective.

This study highlights the value of engineered human multicellular brain tissues for dissecting non-cell-autonomous mechanisms of neurodegeneration. The miBrain platform enables patient-derived modeling, cell-type-specific genetic engineering within a reproducible, cryopreservable brain tissue, capabilities that are difficult to achieve in self-organizing organoids or animal models. Using this platform, we show that *APOE4/4* astrocytes are sufficient to promote pathogenic α-Syn accumulation in miBrains, linking astrocytic cholesterol dysregulation to pathogenic α-Syn propagation.

## Supporting information

KRT

## Limitations of the Study

While miBrains have a user-defined cell composition, some cell types like oligodendroglia progenitors might retain lineage plasticity, resulting in unintended differentiation or proliferation. The *SNCA-A53T* expression system may not fully recapitulate Lewy pathology. Although we identify lipid accumulation and lysosomal dysfunction as central features of *APOE4/4* astrocytes, lipidomic, targeted metabolomic, and organelle-enriched analysis will be required to resolve compartment-specific lipid alterations and their contributions to α-Syn pathogenic conversion. While our results suggest that astrocytes modify α-Syn, the kinases and additional astrocyte-derived factors involved, as well as the persistence of non-astrocytic pathogenic α-Syn remain to be determined. Targeted α-Syn immunodepletion and kinase perturbation studies will help define these mechanisms. Additionally, while microglia and other cell types did not substantially modify the α-Syn phenotypes examined here, they may influence pathological features or treatment responses. Limited microglial recovery in our snRNA-seq dataset, likely due to technical limitations associated with nuclei isolation^110,111^, precluded robust transcriptional analysis. Importantly, our mechanistic conclusions regarding the cell type-specific contributions are based primarily on histological rather than transcriptomic readouts. Continued refinement of the miBrain system, including multicellular functional validation, improved biochemical and transcriptomic workflows, vascular perfusion, and *in vivo* validation of pharmacological treatments, will further expand its utility for modeling human neurodegeneration. Sex-specific effects were not evaluated and remain to be determined.

## Resource availability

### Lead Contact

Further information and requests for resources and reagents should be directed to Dr. Joel W. Blanchard.

### Material availability

This study did not generate new, unique reagents.

## Data and code availability

- The data, protocols, software, and key resources associated with this study are listed in the Key Resources Table (KRT; DOI: 10.5281/zenodo.21177115). The KRT provides links to all deposited datasets, analysis pipelines, and protocols, including raw snRNA-seq sequencing data, processed datasets, and interactive visualization resources.
- Any additional information required to reanalyze the data reported in this paper is available from the lead contact upon request.

## Acknowledgments

We thank Tatyana Kareva for technical support, Natalie Suhy for help with MEA, Anna Bright and Ana Forton-Juarez for help with cellular differentiations, and Dr Elena Coccia for revising the manuscript. We thank Dr Cristina d’Abramo (The Litwin Zucker Center for Alzheimer’s Disease Research) for producing PHF1 antibody and providing guidance on its use. We acknowledge the use of BioRender for generation of cartoons.

This work has been funded in part with federal funds from NASA under contract #80ARC022CA004 titled “Identification of Biomarkers and Pathological Mechanisms via Longitudinal Analysis of Neurological and Cerebrovascular Responses to Neurotoxic Stress Using a Multi-cellular Integrated Model of the Human Brain.” This research was funded in whole or in part by Aligning Science Across Parkinson’s [ASAP-024297] through the Michael J. Fox Foundation for Parkinson’s Research (MJFF). For the purpose of open access, the author has applied a CC BY public copyright license to all Author Accepted Manuscripts arising from this submission. This work was also supported by funding from the NIH/NINDS/NIA (R01NS114239, UH3NS115064, and 1U54AG090669-01), the CureAlz Fund, and the SWT Foundation. C.G. was supported by funding from the NIH/NIA (T32AG04968) and NIH/NINDS (F31NS13090).

## Author Contributions

Conceptualization: L.A.M.L, C.G., and J.W.B.; Methodology: L.A.M.L., C.G., S.G, A.B., D.K, A.H., E.R.S., A.N., J.F.F., E.H., A.U., B.R.S., R.B.R., J.B., J.J.B.C..; Analysis: L.A.M.L., C.G., S.G., A.B., D.K., D.L., J.B., J.J.B.C, P.R..; Writing – Original Draft: L.A.M.L and C.G.; Writing – Review & Editing: L.A.M.L., C.G., J.W.B., A.B., A.H., A.N., D.L., J.B., J.J.B.C., J.F.F, and V.K; Supervision: J.W.B.; Funding Acquisition: J.W.B.

## Declaration of Generative AI and AI-assisted technologies in the writing process

During the preparation of this work, the authors used ChatGPT to improve the grammar and clarity of the manuscript. The technology was not used to generate original scientific content or draft any section of the manuscript from scratch. Following its use, the authors carefully reviewed and edited all text as needed and take full responsibility for the content of the publication.

## STAR★METHODS

### KEY RESOURCES TABLE

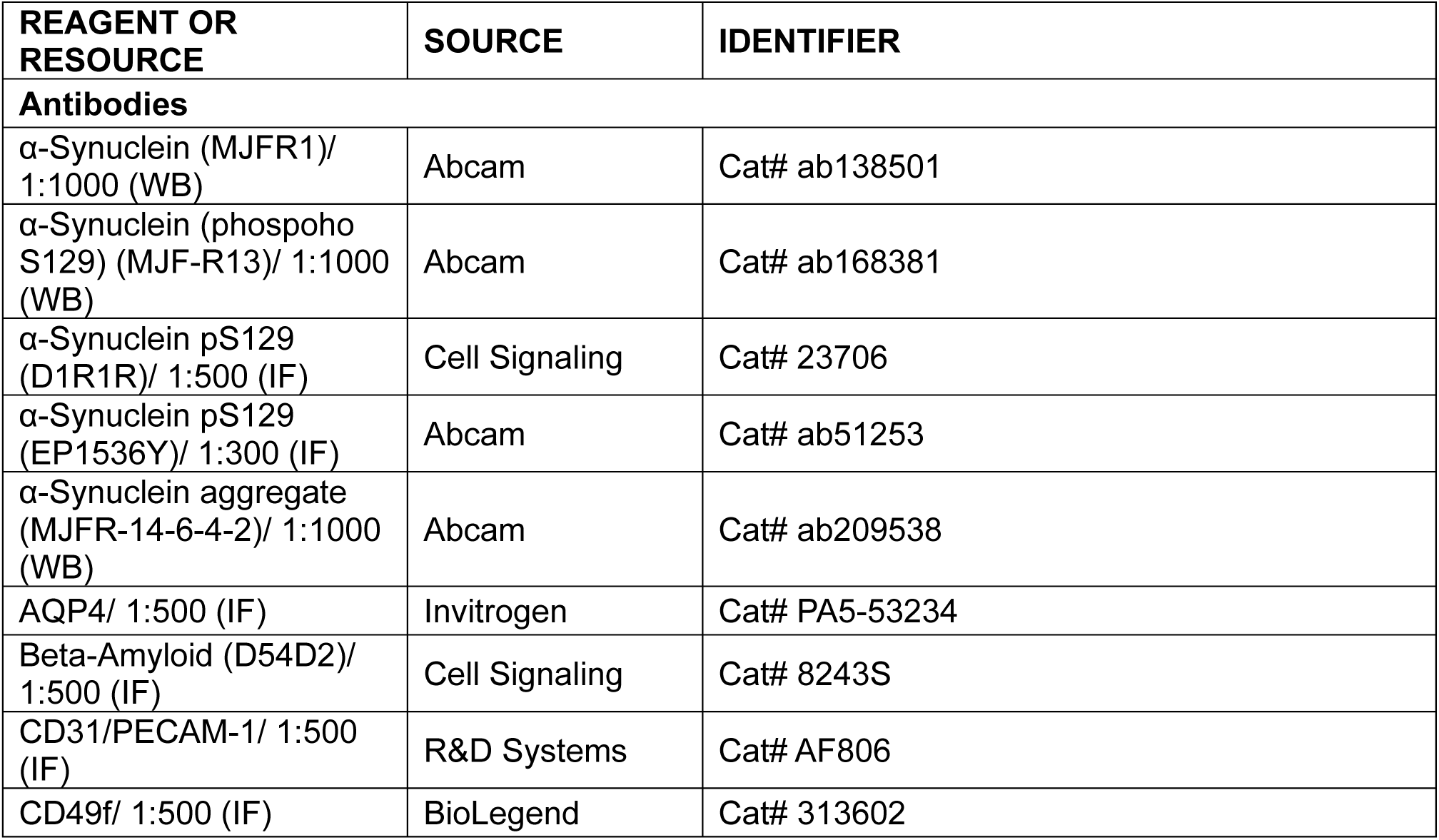
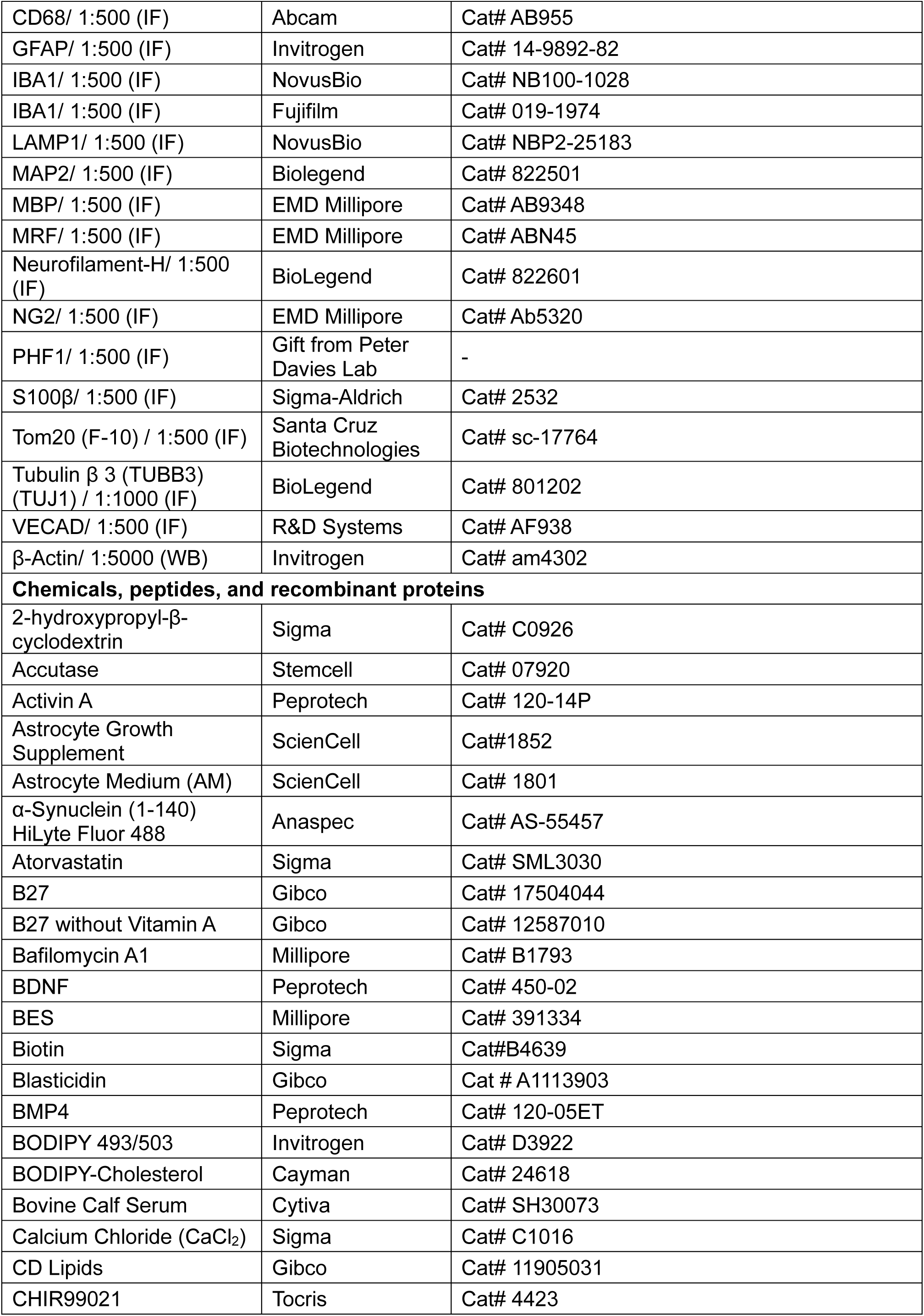
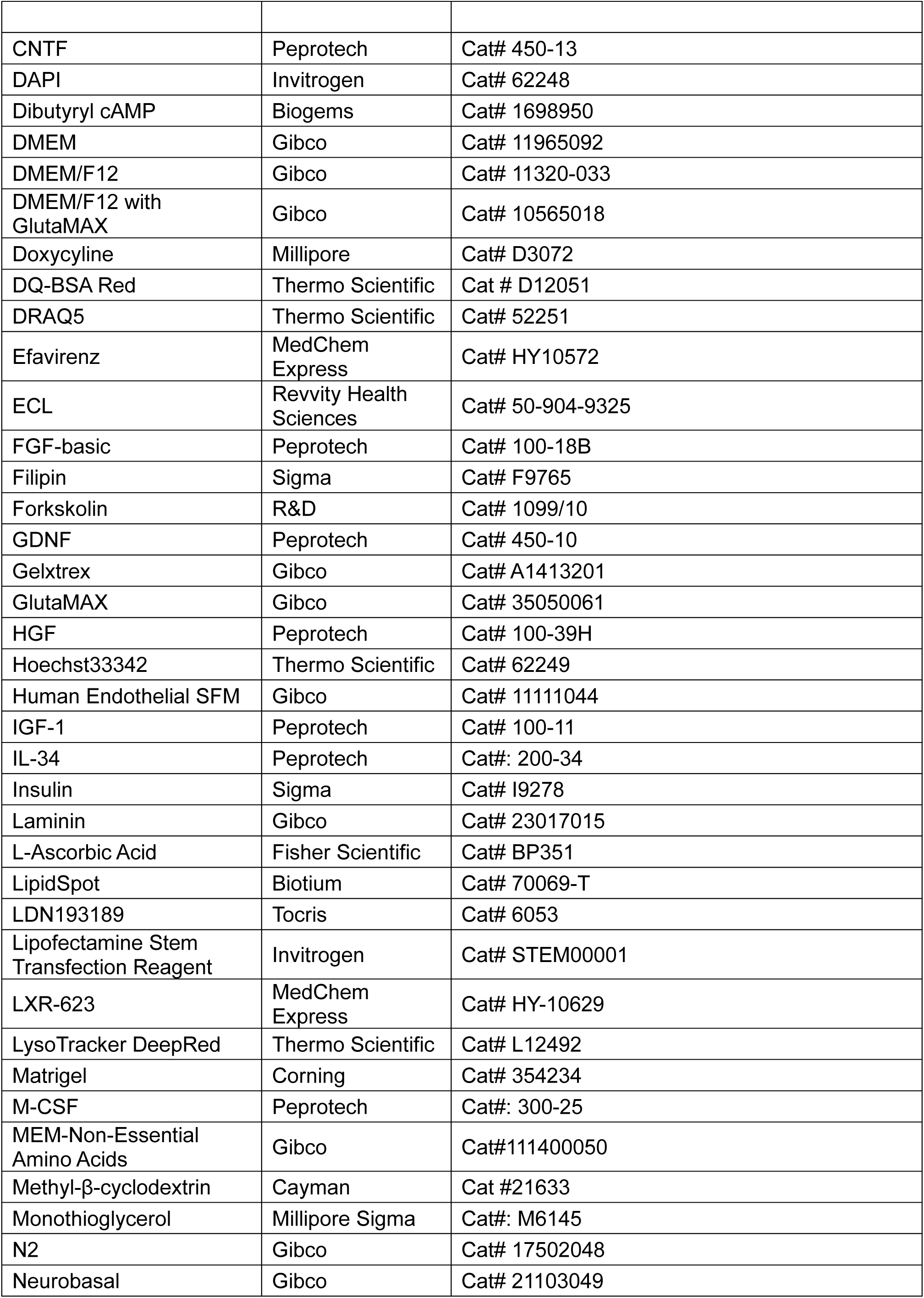
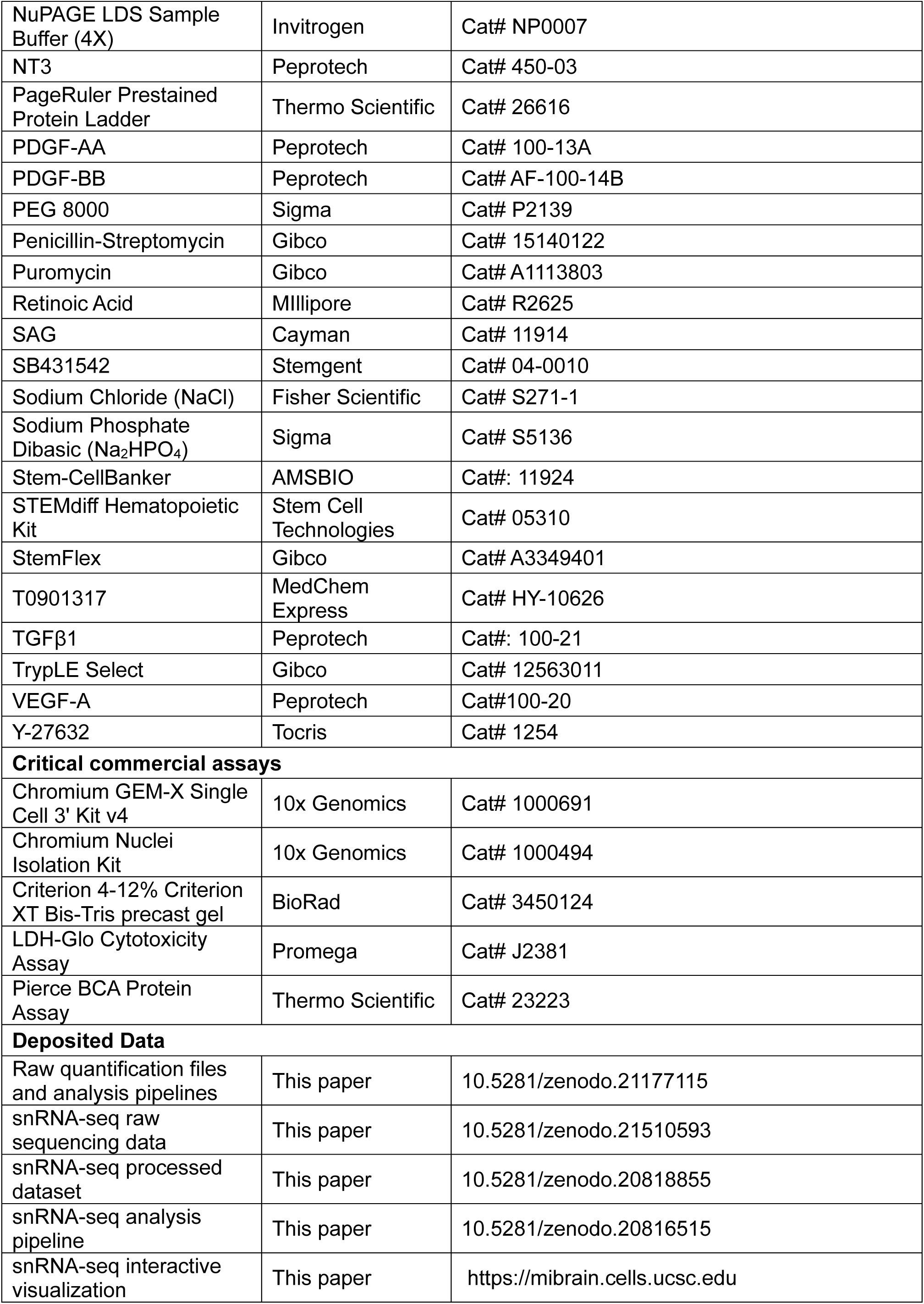
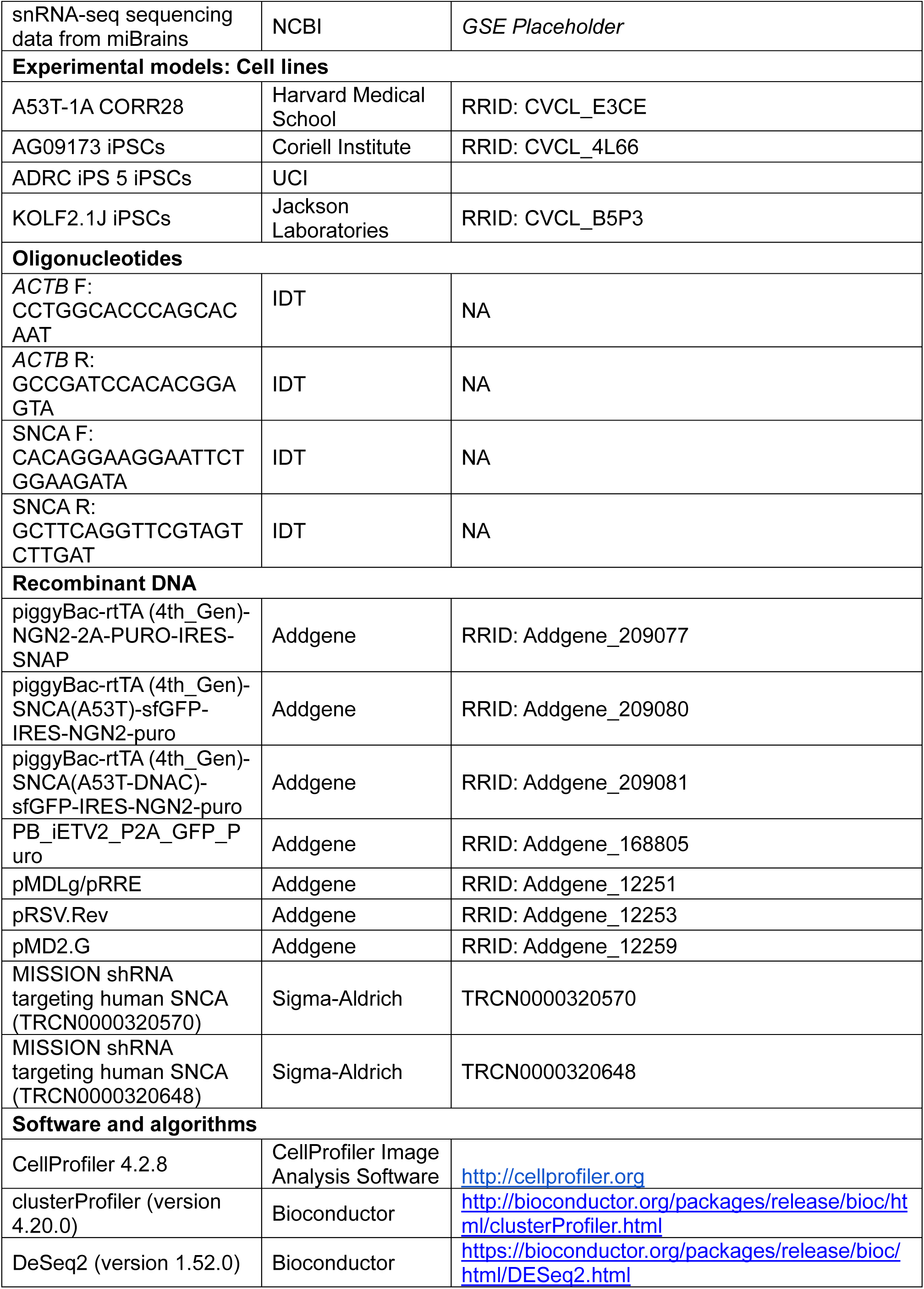
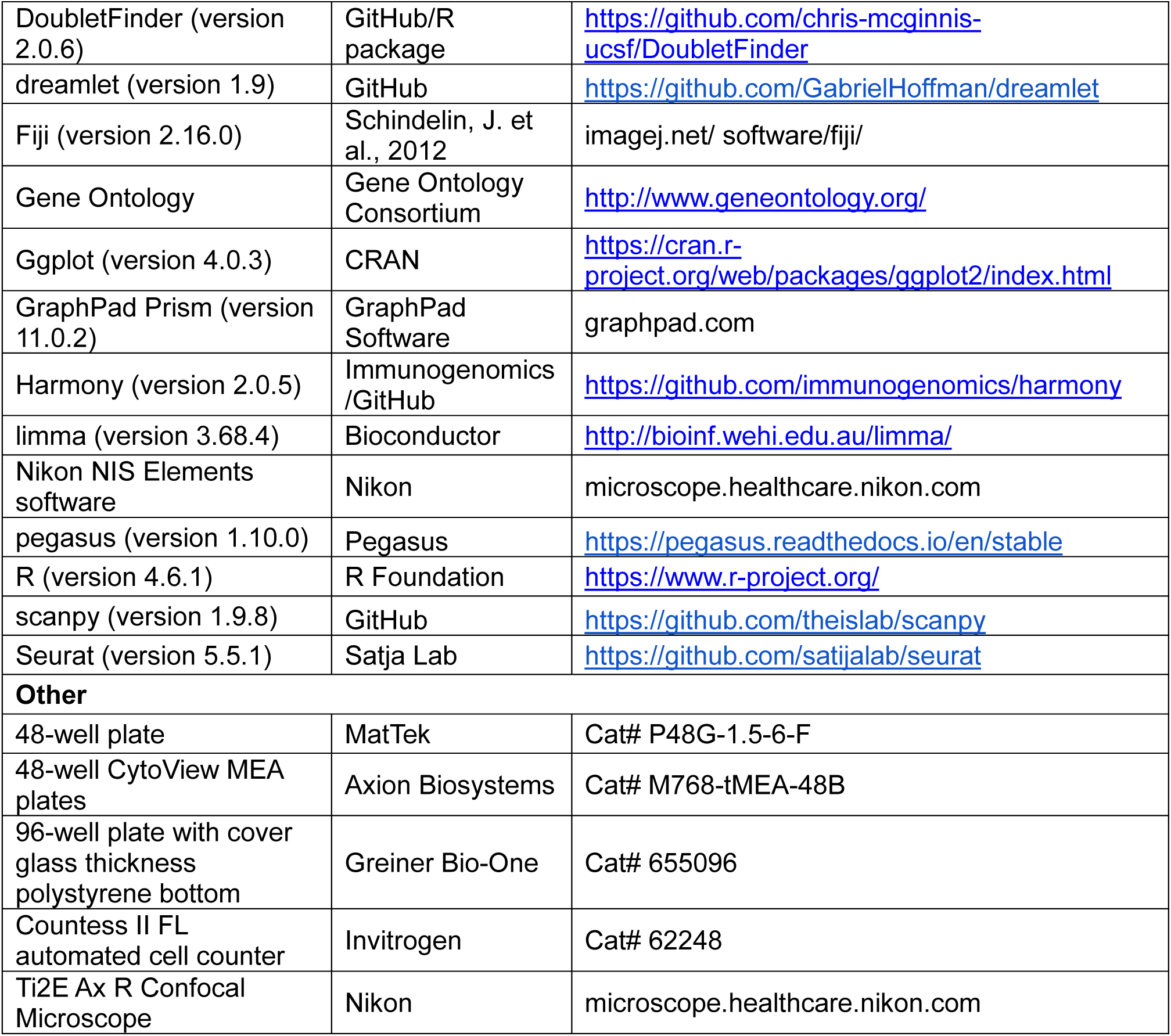

## EXPERIMENTAL MODEL AND STUDY PARTICIPANT DETAILS

### Human iPSC cell lines

All human iPSCs were maintained in feeder-free conditions in StemFlex^TM^ medium (Gibco) on Geltrex^TM^ Matrix (Thermo Fisher Scientific) pre-coated plates. All iPSC lines used in this study are listed in the Key Resource Table. Isogenic *APOE3/3* and *APOE4/4* pairs were generated previously via CRISPR/Cas9^66^. Human iPSCs were grown as colonies in StemFlex^TM^ medium until they reached 60-70% confluency. At this point, iPSCs were either passaged for maintenance using 0.5 mM EDTA to gently lift colonies or harvested using Accutase^TM^ cell detachment solution for 5-10 minutes at 37°C to start a differentiation protocol as singularized cells. A detailed iPSC maintenance protocol is available at doi.org/10.17504/protocols.io.bp2l6xmjrlqe/v1. A total of four donor cell lines were used in this study (see Key Resources Table). For all figures where the donor number is not indicated, we have used Donor 1 (AG09173 iPSCs). Donor 2 corresponds to KOLF2.1J iPSCs (*APOE3/3* parental: JIPSC001000; *APOE4/4* Clone A: JIPSC001150) and Donor 3 to ADRC iPS 5 iPSCs. The cell line A53T-1A CORR28 was obtained from a patient harboring the SNCA A53>T mutation, which was mutation-corrected, and then received the plasmid for overexpression of SNCA-A53>T. This strategy was used to introduce the mutation on SNCA at overexpressed levels while keeping the rest of the patient’s “synucleinopathy-permissive” genetic background. This line was kindly provided by Dr. Vikhram Khurana after extensive characterization^29,113^, and its use is indicated in the methods section and figure legends. Prior to all experiments, the APOE genotype of each cell line was confirmed by Sanger sequencing, and only lines with the verified *APOE3/3* or *APOE4/4* genotype were used.

## METHOD DETAILS

### Differentiation of human iPSCs into neurons

Neuron differentiation was adapted from Zhang *et al*.^28^ and Lam *et al*.^29^. Briefly, iPSCs were transfected with PiggyBac plasmids to confer doxycycline-inducible expression of the Neurogenin-2 gene (*NGN2*, Addgene Plasmid #209077) alone or in combination with SNCA-A53T-sfGFP (Addgene Plasmid: 209080) or A53T-ΔNAC-SNCA-sfGFP (Addgene Plasmid: 209081), using Lipofectamine^TM^ Stem Transfection Reagent. Briefly, dissociated iPSCs were plated at ∼104,000 cells/cm^2^ onto Geltrex^TM^-coated plates, in StemFlex™ supplemented with 10 µM Y27632 and 5 µg/mL doxycycline (day 0). At day 1, medium was replaced with Neurobasal N2B27 medium (Neurobasal, 1x B-27, 1x N-2, 1x MEM-NEAA, 1x GlutaMAX, 1% penicillin-streptomycin) supplemented with 10 µM SB431542, 100 nM LDN, 5 µg/mL doxycycline, and 5 µg/mL Blasticidin. On day 2, the medium was replaced with Neurobasal N2B27 media supplemented with 10 µM SB431542, 100 nM LDN, 5 µg/mL doxycycline, and 1 µg/mL puromycin. On days 3-6, the medium was replaced daily with Neurobasal N2B27 media supplemented with 5 µg/mL doxycycline and 1 µg/mL puromycin. At day 7, cells were dissociated with Accutase and either seeded into miBrains (see below) or seeded into 2D monocultures on Poly-L-Ornithine and laminin pre-coated plates at 156,250 cells/cm^2^, in Neurobasal N2B27 with 5 µg/mL doxycycline and 10 µM Y27632. For 2D cultures, on day 8, wells were gently topped with Neurobasal N2B27 supplemented with 20 ng/mL BDNF, 20 ng/mL GDNF, 1 mM dcAMP, 2 µg/mL laminin, and 1 µM AraC, using the same volume of medium in the wells. At day 11, media is replaced with Neurobasal N2B27 supplemented with 10 ng/mL BDNF, 10 ng/mL GDNF, 0.5 mM dcAMP, and 1 µg/mL laminin. This medium was used for half-media changes every 3-4 days.

### Differentiation of human iPSCs into astrocytes

Astrocytes were generated using previously published protocols for iPSC-derived NPC (Chambers *et al*.)^30^ and astrocyte (TCW *et al*.)^31^ differentiations. Briefly, dissociated iPSCs were plated at 100,000 cells/cm^2^ onto Geltrex^TM^-coated plates, in pre-warmed StemFlex™ supplemented with 10 µM Y27632. Cells were fed every other day with StemFlex™ until they reached >95% confluence. Once cells reached confluence, the medium was replaced with NPC medium (1:1 DMEM/F12: Neurobasal Medium, 1x N-2 Supplement, 1x B-27 Serum-Free supplement, 1x GlutaMAX Supplement, 1x MEM-NEAA, 1% penicillin-streptomycin) supplemented with 10 µM SB43152 and 100 nM LDN193189 (day 0). From days 1 to 9, cells were fed daily with NPC medium plus 10 µM SB43152 and 100 nM LDN193189. At day 10, cells were split with Accutase and replated onto fresh Geltrex^TM^-coated plates, in NPC media supplemented with 20 ng/mL bFGF and 10 µM Y27632. From days 11 to 13, cells were fed with NPC media plus 20 ng/mL bFGF. At day 14, cells were split with Accutase and re-seed onto fresh Geltrex^TM^-coated plates, in NPC media plus 20 ng/mL bFGF and 10 µM Y27632. Starting from day 15, cells were fed every 2-3 days with Astrocyte Medium (AM, ScienCell) and passaged using Accutase once they reached 90% confluence. From this point, NPCs were fully differentiated into astrocytes in 30 days. NPCs and fully differentiated astrocytes were cryopreserved in a freezing medium consisting of 90% knockout serum replacement (KSR) and 10% dimethyl sulfoxide (DMSO). A detailed protocol is available at doi.org/ 10.3791/65921.

### Differentiation of human iPSC into brain microvascular endothelial cells

Brain endothelial cell differentiation was adapted from the protocols from Blanchard *et al.*, Qian *et al.,* and Wang *et al*. ^32,35,114^. Briefly, iPSCs were transfected with a PiggyBac plasmid to confer doxycycline-inducible expression of the ETS variant transcription factor 2 (*ETV2*, Addgene Plasmid #168805), using Lipofectamine^TM^ Stem Transfection Reagent. Inducible ETV2-iPSCs were grown until 60-70% confluency, dissociated with Accutase, and plated at 20,800 cells/cm^2^ onto Geltrex^TM^-coated plates in StemFlex™ supplemented with 10 µM Y27632 (day 0). On day 1, the medium was replaced with DeSR1 medium (DMEM/F12 with GlutaMAX, 1× MEM-NEAA, 1× penicillin-streptomycin) supplemented with 10 ng/mL BMP4, 6 µM CHIR99021, and 5 µg/mL doxycycline. On day 3, the medium was replaced with DeSR2 medium (DeSR1 media, 1x N-2, 1× B-27) supplemented with 5 µg/mL doxycycline. At days 5 and 7, the medium was replaced with hECSR medium (Human Endothelial Serum-free Medium, Gibco, 1× MEM-NEAA, 1× B-27, 1% penicillin-streptomycin) supplemented with 50 ng/mL VEGF-A, 2 μM Forskolin, and 5 µg/mL doxycycline. At day 8, cells were dissociated using Accutase and re-seed onto fresh Geltrex^TM^-coated plates in hECSR supplemented with 50 ng/mL VEGF-A and 5 µg/mL doxycycline. This medium was used for every 2-3 days medium change to maintain cells for up to 1 week until ready for tissue assembly (miBrain or JAMs). A detailed protocol is available at doi.org/10.1007/978-1-0716-3287-1_11.

### Differentiation of human iPSCs into mural cells

Mural cells were differentiated using previously published protocol from Patsch *et al.*^36^. Dissociated iPSCs were plated at 37,000 to 40,000 cells/cm^2^ onto Geltrex^TM^-coated plates, in StemFlex™ supplemented with 10 µM Y27632 (day 0). On day 1, the medium was replaced with N2B27 medium (1:1 DMEM/F12: Neurobasal media, 1x B-27, 1x N-2, 1x MEM-NEAA, 1x GlutaMAX, 1% penicillin-streptomycin) supplemented with 25 ng/mL BMP4 and 8 μM CHIR99021. At days 3 and 4, the medium was replaced with N2B27 media supplemented with 10 ng/mL Activin A and 10 ng/mL PDGF-BB. At day 5, mural cells were dissociated with Accutase and re-seeded onto fresh 0.1% gelatin-coated plates at 35,000 cells/cm^2^, in N2B27 supplemented with 10 ng/mL PDGF-BB. This medium was used every 2-3 days medium was change for another 5–7 days. Cells were then banked in freezing medium (90% KSR/ 10% DMSO) and expanded in N2B27 until ready for tissue assembly (miBrain or JAMs). A detailed protocol is available at doi.org/10.1007/978-1-0716-3287-1_11.

### Differentiation of human iPSCs into oligodendrocyte progenitor cells (OPCs)

OPC differentiation was adapted from Douvaras et al, 2014^115^. Briefly, iPSCs were dissociated into single cells using Accutase and seeded at near-confluent density. Differentiation began the next day (designated as day 0) by culturing the cells in DMEM/F12 (1:1) medium supplemented with N2, 10 μM SB431542, 100 nM LDN 193189, and 100 nM all-trans retinoic acid (RA), with daily medium changes. On day 8, 1 μM SAG was added to the differentiation medium. By day 12, adherent cells were detached and transferred to low-attachment plates to form cell spheres. These spheres were cultured in DMEM/F12 (1:1) medium containing N2, RA, and SAG. On day 30, spheres were plated onto poly-L-ornithine/laminin-coated plates to allow cells to migrate outward. At this stage, the medium was replaced with DMEM/F12 (1:1) supplemented with N2, B27, 10 ng/ml PDGF-AA, 10 ng/ml IGF, 5 ng/ml HGF, 10 ng/ml NT3, 25 μg/ml insulin, 100 ng/ml biotin, 1 μM cAMP, and 60 ng/ml T3. By day 75, cells were harvested, dissociated, and purified using NG2-specific magnetic-activated cell sorting (MACS). The enriched cells were expanded in DMEM/F12 (1:1) medium supplemented with N2, B27 without Vitamin A, 10 ng/ml PDGF-AA, 10 ng/ml β-FGF, and 10 ng/ml NT3 until ready for tissue assembly (miBrain or JAMs).

### Differentiation of human iPSCs into microglia

iPSC-derived microglia were differentiated as previously shown^38,116^ via an intermediate differentiation step into hematopoietic progenitor cells (HPCs). For the generation of HPCs, the STEMdiff Hematopoietic Kit was used, according to the manufacturer’s manual. Briefly, when 70% confluent (day 0), iPSCs were harvested and passaged at a density of 20-40 colonies per well in a 6-well coated with 0.1mg/mL Matrigel. On day 1, Medium A was added to the culture, and on day 4, it was switched to Medium B until complete HPC maturation on days 11-13. Fully differentiated HPCs, floating in the medium and detached from the colonies, were collected for microglial differentiation or frozen in Stem-CellBanker supplemented with microglial cytokines. For the generation of mature microglia, differentiated HPCs were collected and transferred into Matrigel coated 6-well plate at a confluency of 350k HPCs per well. The differentiation takes 25-28 days, during which HPCs are cultured in microglia media consisting of DMEM/F12, with 2X B27, 0.5X N2, 1X Glutamax, 1X non-essential amino acids, 400 mM Monothioglycerol, and 5 ug/mL human insulin, freshly supplemented with 100 ng/mL IL-34, 50 ng/mL TGFβ1, and 25 ng/mL M-CSF. Microglia were added to miBrains within the pool of Geltrex encapsulated cells at the time of miBrain assembly. MiBrains were maintained in miBrain media supplemented with 100 ng/mL IL-34 and 25 ng/mL M-CSF for 1 week and then switched to miBrain media supplemented with 25 ng/mL M-CSF until downstream experiments.

### 3D Tissue Assembly for miBrains

Neurons, astrocytes, endothelial cells, mural cells, and OPCs were dissociated using Accutase or TryplE Select (astrocytes). Cells were resuspended in corresponding media, counted, and resuspended at 1 x 10^6^ cells/ mL. For miBrains, a tube was prepared to contain 5 x 10^6^ neurons, 5 x 10^6^ endothelial cells, 1 x 10^6^ astrocytes, 1 x 10^6^ OPCs, and 1 x 10^6^ mural cells per 1mL of miBrain. Microglia were added for a subset of miBrains (indicated in the figure legends) at a ratio of 1.67 x 10^6^ per 1 mL. Pooled cells were spun down at 200 x g for 5 minutes at RT. Media was aspirated carefully, leaving the cell pellet undisturbed. The cell pellet was placed on ice and resuspended in 1 mL Geltrex^TM^ supplemented with 10 µM Y27632 and 5 µg/mL doxycycline, avoiding air bubbles and keeping it on ice to prevent premature Geltrex polymerization and inability to seed miBrains properly. To generate miBrain tissue that adopted a free-floating, organoid-like morphology over time, 25-50 µL of encapsulated cell suspension were seeded per inner glass-bottom well of a 48-well MatTek plate (MatTek). To generate miBrain tissue that remained attached to the plate (more suitable for automated imaging) while conserving 3D morphology, 10 µL of encapsulated cell suspension were seeded per well of a 96-well µClear plastic-bottom plate (Greiner). After miBrains were seeded, the plates were transferred into a 37 °C 95%/5% Air/CO2 incubator for 30 minutes to allow the Geltrex^TM^ to polymerize. After polymerization of the gel, miBrain week-1 medium (Human Endothelial Serum-free Medium, 1x Pen/Strep, 1X MEM-NEAA, 1X CD Lipids, 1x Astrocyte Growth Supplement (ScienCell), 1x B27 Supplement, 10µg/mL Insulin, 1µM cAMP-dibutyl, 50µg/mL ascorbic acid, 10ng/mL NT3, 10ng/mL IGF, 100ng/mL biotin, 60 ng/mL T3, 50 ng/mL VEGF, 1µM SAG, 5 µg/mL doxycycline) was added to each well, ensuring complete submersion of the culture in media (250-500uL per well of a 48-well matTek plate, 100 to 200 uL per well of a 96-well plate). Half media was changed every 2-3 days. On day 8 after miBrain seeding, the media was changed to miBrain week-2 medium (Human Endothelial Serum-free Medium, 1x Pen/Strep, 1X MEM-NEAA, 1X CD Lipids, 1x Astrocyte Growth Supplement (ScienCell), 1x B27 Supplement, 10µg/mL Insulin, 1µM cAMP-dibutyl, 50µg/mL ascorbic acid, 10ng/mL NT3, 10ng/mL IGF, 100ng/mL biotin, 60 ng/mL T3, 5 µg/mL doxycycline). Cultures were used for downstream assays after 18 days, unless otherwise indicated.

### Cryopreservation of miBrains

For JAMs assembly and cryopreservation, a tube containing 5 x 10^6^ endothelial cells, 1 x 10^6^ astrocytes, 1 x 10^6^ OPCs, and 1 x 10^6^ mural cells per 1 mL was prepared. Pooled cells were spun down at 200 x g for 5 minutes at RT and cryopreserved in miBrain freezing media (60% KSR, 30% hECSR medium, 10% DMSO, 10 µM Y27632, 50 ng/mL VEGF-A). Upon thaw, cell viability was assessed, and the appropriate volume of neurons needed to conserve the original miBrain cell-to-cell ratio was added to the pooled cell suspension, which was spun again, encapsulated, and seeded as described above.

### Basal electrophysiological activity in miBrains

To functionally validate the neuronal networks within miBrains, we performed microelectrode array (MEA) recordings under basal conditions. Basal electrophysiological activity of miBrains was assessed, as described previously^117^, with minor modifications, using 48-well CytoView MEA plates containing 16 electrodes per well. Briefly, plates were pre-coated with 0.1% polyethylenimine in borate buffer, rinsed, air-dried overnight, and coated with 20 µg/ml laminin immediately before use. Individual miBrains were transferred to coated MEA wells and allowed to settle for 24 h before recording. Spontaneous extracellular activity was recorded at 37 °C using a Maestro Pro MEA system and AxIS Spontaneous Neural configuration. Spikes were detected using an adaptive threshold set to 5X the standard deviation of the estimated noise per electrode. Electrodes detecting ≥5 spikes/min were classified as active, and network bursts were defined as events containing ≥5 spikes with inter-spike intervals <100 ms occurring in at least 20% of active electrodes; synchrony index was calculated using a 20 ms cross-correlogram window. Mean firing rate, burst parameters, network burst frequency, and synchrony index, among other parameters, were quantified using Neural Metric Tool (Axion Biosystems), and brightfield images were acquired to assess electrode coverage by miBrains organoids.

### Exposure of tissue to exogenous α-Synuclein

miBrains were exposed to 4 µg/ml of Human Recombinant Alpha Synuclein Protein Aggregates (Pre Formed Fibrils, PFFs, from StressMarq, #SPR-322, or Abcam, #ab218819) or 4 µg/mL patient-derived PFFs (gift from Vik Khurana). PFFs were sonicated in a water bath (VWR Ultrasonic Cleaner) immediately before adding to the cultures, using 10 cycles of 30 seconds on and 30 seconds off. Free-floating miBrains incubated with α-Syn had a media change after 48 hours and were fixed in 4% paraformaldehyde after 2 weeks.

### Induction of α-Syn pathology via expression of SNCA-A53T

Neurons with inducible expression of *SNCA* with the A53>T mutation (A53T) were generated from iPSC as described above. Cells were harvested on day 7 of the differentiation and cultured in 2D on Poly-L-Ornithine and laminin pre-coated plates at 156,250 cells/cm^2^,or encapsulated for 3D cultures in Geltrex™ at 50,000 cells for 10 µL of Geltrex™, or added to JAMs for miBrain assembly as described above. The cultures were maintained for 18 days with or without PFFs. For the cultures that received PFFs, 4 µg/ml PFFs (gift from Vik Khurana) were added to the media on day 4 after assembly. The media was half-changed every 2-3 days.

### Immunofluorescence

2D cultures were fixed in 4% paraformaldehyde for 15 minutes at room temperature and rinsed with PBS. 2D cultures (non-neuronal) were blocked in 0.3% Triton-X100, 5% normal donkey serum in PBS for 30 minutes., and then with primary antibodies diluted in blocking buffer overnight at 4°C. Cultures were rinsed 3 times with 0.3% Triton-X100 in PBS for 15 minutes each and incubated with secondary antibodies (and Hoechst 33342) diluted at 1:1000 in blocking buffer for 2 hours at room temperature. For neutral lipid and cholesterol staining, BODIPY 493/503 and filipin were added at 2μM and 0.1/mg/mL respectively in PBS for 1 hour after secondary incubation. Cultures were rinsed 3 times with PBS for 15 minutes and left in PBS for image acquisition. 2D neuronal cultures were blocked in 0.1% Triton-X100, and 10% normal donkey serum in PBS, antibodies were diluted in 0.02% Triton-X100 and 2% normal donkey serum in PBS, and washes were done with PBS. All other conditions were kept the same as the other cell types.

3D cultures and miBrains were fixed in 4% paraformaldehyde overnight at 4°C and rinsed with PBS. 3D cultures and miBrains were incubated in blocking solution (0.3% Triton-X100, 5% normal donkey serum, in PBS) overnight at 4°C and then with primary antibodies diluted in blocking solution for 2-3 nights at 4°C. All antibodies were diluted at 1:500 except for α-Synuclein pS129 (EP1536Y) which was diluted at 1:300. Cultures were rinsed 5 times with 0.3% Triton-X100 PBS for 30 minutes each and incubated with secondary antibodies and Hoechst 33342 (Thermo 62249) diluted at 1:1000 in blocking solution for 2-3 nights at 4°C. Cultures were rinsed 5 times with 0.3% Triton-X100 PBS for 30 minutes each then rinsed and left in PBS for image acquisition.

### Image acquisition and quantification

Images were acquired using a confocal microscope (Leica Stellaris, Nikon AX R, or Thermo Scientific CellInsight CX7). For quantification, single optical sections or 10-20 mm Z-stack images were acquired at 10x or 20x, 4-8 fields per sample. Area, Intensity or Volume measurements on single images or Z-stacks were performed using Nikon AX R built-in quantification software, ImageJ, or CellProfiler. All area and volume analysis were performed using automated pipelines including the application of the same image threshold for all experimental conditions in the same experiment.

### Preparation of miBrains for single-nucleus RNA sequencing (snRNA-seq)

miBrains were cultured in standard conditions for 18 days, rinsed with PBS and frozen without media. Fresh frozen miBrains were processed using the Chromium Nuclei Isolation Kit (10x Genomics, #1000494), with RNase inhibitors. Nuclei were quantified via DAPI (Invitrogen, #62248) staining using a Countess II FL automated cell counter (Invitrogen, #AMQAF1000). 20,000 nuclei per sample were processed using 3’ GemX v4 reagents (10x Genomics, #1000691), and libraries prepared according to the manufacturer’s instructions (10xgenomics.com). All libraries were sequenced at Azenta using the NovaSeq X platform (Illumina).

### miBrain snRNA-seq Data Processing

Raw sequencing reads were aligned to the human reference genome (GRCh38) and per-nucleus count matrices generated using Cell Ranger (10x Genomics, v9.0.1). Six libraries were processed — two replicates each for Control, *APOE3/3-SNCA-*A53T, and *APOE4/4-SNCA-*A53T conditions. Nuclei were filtered out if they had unique molecular identifier (UMI) counts or number of detected genes outside ±4 median absolute deviations (MADs) from the sample median, or if the fraction of mitochondrial reads exceeded 5%. Doublets were identified and removed using the simulation-based inference implemented in pegasus v1.10.0^118^, stratified by sample, prior to final clustering. Library-size normalization followed by log(x+1) transformation was then applied using scanpy v1.9.8^119^. Highly variable genes (HVGs) were identified using a Cell Ranger-flavored dispersion method using scanpy^119^, computed within each sample independently to account for inter-library variability, and restricted to protein-coding autosomal genes, excluding ribosomal (RPL/RPS), mitochondrial, and MitoCarta-annotated genes^120^. Principal component analysis (PCA) was performed on the 6,000 HVG-filtered, normalized matrix using pegasus v1.10.0, retaining 30 informative PCs identified from an elbow plot. Technical covariates (total UMI count, mitochondrial fraction, and cell cycle score) were regressed from the PCA embedding. Cross-sample batch effects were corrected using Harmony^121^. A k-nearest neighbor graph was constructed on the corrected embedding (K = 100) and Leiden community detection (resolution =0.2)^122^. UMAP embeddings were computed on the harmonized PCA space.

To annotate the clusters, we first identified the most specific genes in each cluster using the cellTypeSpecificity function in dreamlet v1.9^123^. We then calculated the Spearman correlation between the average cluster-level expression of cell-type-specific genes and the annotated related cell types from Tereza Clarence et al.^112^. Cell-level metadata, including QC metrics and cell-type annotations, are provided in **Table S3**. QC metrics per sample are available in **Table S4**. Annotated cell numbers per sample are available in **Table S5**.

Following cell type annotation, for pseudo-bulk differential expression, raw counts were aggregated per cell type and sample using aggregateToPseudoBulk from dreamlet v1.9^123^, producing one pseudo-bulk profile per cell type–sample combination. Within each cell type, counts were normalized and precision weights estimated using the voom method via processAssays (minimum count per gene = 5). Pairwise differential expression between conditions was performed per cell type using linear mixed models implemented in dreamlet, with condition as the fixed effect and number of detected features as a covariate.

To test whether biologically focused gene programs were transcriptionally dysregulated in *A53T APOE3/3* excitatory neurons compared to controls and in *APOE4/4* astrocytes compared to *APOE3/3*, we performed gene set enrichment analysis (GSEA). We used manually curated gene sets directly relevant to Parkinson’s disease and lipid biology (**Table S2**). Genes were ranked by their t-statistic from the dreamlet pseudo-bulk linear model (coefficient: *A53T-APOE3* vs. control, neurons; *A53T-APOE4* vs. *A53T-APOE3*, astrocytes). We then ran GSEA on these custom gene sets using clusterProfiler v4.8.1^124^. In parallel, we tested the same ranked gene list against Gene Ontology (GO) Biological Process relevant to protein phosphorylation regulation (protein dephosphorylation, GO:0006470).

### Human iPSC-derived astrocyte bulk RNAseq analysis

Fragments Per Kilobase of transcript per Million mapped reads (FPKM) values were obtained from published bulk RNA-seq data of isogenic human iPSC-derived astrocytes expressing either *APOE3/3* or *APOE4/4* (GEO: GSE102956)^66^. FPKM values were log2-transformed with an offset of 0.1 and genes with zero variance across all the samples were excluded. To focus on genes with meaningful variability, an additional filtering step was applied to retain genes above the 10th percentile (variance > 0.033) after the removal of the zero-variance genes. Differential expression analysis was performed using the limma package in R. A design matrix was constructed to model the experimental conditions (APOE3 vs. APOE4) and sample-specific array weights were estimated using the array Weights function with a prior.n value of 100 to help stabilize weight estimation. A linear model was then fit to the log-transformed, filtered expression data using lmFit, and empirical Bayes moderation was applied via the eBayes function, with trend and robust both set to TRUE. DEGs were defined as having an absolute log2 fold change > 0.5 and adjusted p-value (false discovery weight) < 0.05. A volcano plot was then generated using the ggplot2 package and genes meeting the DEG thresholds were highlighted. Enrichment analysis was performed on the upregulated and downregulated DEGs separately using the clusterProfiler package. Pathways were identified through Gene Ontology (GO) database. The top 10 pathways for each direction (upregulated and downregulated) were selected based on adjusted p-values.

### Human astrocyte scRNAseq pseudobulk analysis

The processed dataset was downloaded from Haney et al (GEO: GSE254205).^73^. Unless otherwise stated, all following analyses were performed in R using the package Seurat^125^. Aligning with the quality control protocol from the source publication, cells with nFeature < 500, nCount < 1000, and mitochondrial and ribosomal reads > 10% were discarded. Doublets were removed using the DoubletFinder package^126^. Following the standard Seurat pipeline with default parameters, raw gene counts were normalized, the top 2,500 highly variable genes were identified, and the data were scaled. Harmony was used to integrate the patient samples, and the top 20 principal components were used in Seurat’s FindNeighbors, FindClusters (0.2 resolution), and RunUMAP functions. Cell-type annotation was manually performed using the marker genes described in Haney et al., and clusters not enriched for these marker genes were removed from further analysis. Pseudobulked samples were generated by summing the raw counts for each patient sample and normalized using Seurat’s AggregateExpression() function.

### Isogenic iPSC-derived astrocyte RNAseq analysis

The raw counts from iPSC-derived astrocyte bulk RNAseq were downloaded from Lin et al (GEO: GSE102956).^66^ Raw counts were processed using the DeSeq2 package^127^ and normalized using the counts() function with normalized set to TRUE.

### DQ-BSA proteolytic activity assay

Astrocytes were seeded at 10,000 cells/well of a 96 well µClear plastic bottom plate (Greiner 655090) in AM. The following day, the media was changed to FBS-free maturation media (50% DMEM/F12, 50% Neurobasal, 1X B27 without Vitamin A, 1X N2, 1X NEAA, 1X GlutaMAX, 10ng/mL CNTF, 1% penicillin-streptomycin). Half of the wells were treated with 100 nM BafilomycinA1 to inhibit lysosomal function as a negative control. The next day, cells were pulsed with 1µg/mL DQ-BSA (Thermo D12051) in maturation media for 30 minutes. Media was replaced with fresh media (and fresh bafilomycin in the appropriate wells), and the red fluorescence and bright field were imaged at 20x with an Incucyte (Sartorius) every hour for 24 hours.

Alternatively, astrocytes were seeded at 50K cells/well of a 12-well plate. Media changes were followed as above. Cells were lifted to single-cell suspension with TrypLE Select 24 hours after the DQ-BSA pulse and filtered into flow cytometry tubes with DAPI. Red fluorescence in live cells was analyzed with a BD Celesta flow cytometer.

### Live Imaging of α-Syn-HiLyte Uptake and Degradation

Astrocytes were seeded 10,000 cells/well of a 96-well µClear plastic bottom plate (Greiner) in AM. The following day, the media was changed to FBS-free maturation media (50% DMEM/F12, 50% Neurobasal, 1X B27 without Vitamin A, 1X N2, 1X NEAA, 1X GlutaMAX, 10ng/mL CNTF, 1% penicillin-streptomycin) and cultured for 2 more days. α-Syn-HiLyte (AnaSpec AS-55457) was added to the media at a final concentration of 2 µg/mL, and nuclei were stained with 10 µg/mL Hoechst 33342. Images were acquired every 30-90 minutes for the first 6 hours on a Nikon AX R. At 24 h, the media was changed to fresh FBS-free maturation media to remove any excess α-Syn. Images were acquired at 24 hours and 48 hours using the same parameters and laser settings.

### Live Imaging of BODIPY-Cholesterol

Astrocytes were seeded 10,000 cells/well of a 96-well µClear plastic bottom plate (Greiner) in AM. The following day, the media was changed to FBS-free maturation media (50% DMEM/F12, 50% Neurobasal, 1X B27 without Vitamin A, 1X N2, 1X NEAA, 1X GlutaMAX, 10ng/mL CNTF, 1% penicillin-streptomycin) and cultured for 3 more days. Astrocytes were incubated with 2 µM BODIPY-Cholesterol (Cayman 24618) for 2 hours. Cells were then incubated with 10 µg/mL Hoechst 33342 for 10 minutes to stain nuclei. Media was replaced with fresh media and cells were imaged using a CX7 High Content Screening Platform with a 20x objective lens (Thermo; LUCPLFLN20x).

### Western blotting and dot blotting

Sequential detergent extraction was performed to biochemically separate detergent-soluble and detergent-insoluble α-synuclein species, adapted from Lam et al., 2024^29^. Briefly, cells lysed in 1% (vol/vol) Triton X-100 in TBS supplemented with protease and phosphatase inhibitors. Lysates were subjected to water bath sonication (VWR Symphony Ultrasonic Cleaner; 30 s on/30 s off, 5 cycles), incubated on ice for 30 min, and centrifuged at 15,000 × g for 30 minutes at 4 °C. The resulting supernatant was collected as the Triton X-100–soluble fraction, and protein concentration was determined by BCA assay. The remaining pellet was resuspended in Triton X-100 buffer, re-sonicated, and centrifuged again at 15,000 × g for 30 minutes. The insoluble pellet was then resuspended in 2% (vol/vol) SDS in TBS containing protease and phosphatase inhibitors and subjected to additional water bath sonication (30 s on/30 s off, 5 cycles) to generate the SDS-soluble fraction. The SDS-soluble supernatant was collected and adjusted to twice the volume of the matched Triton X-100 fraction to enrich for detergent-insoluble α-synuclein species and facilitate visualization on immunoblots. Samples were heated at 65 °C for 5 minutes prior to gel loading. Equal volumes of Triton X-100–soluble and SDS-soluble fractions (20 µL per lane, corresponding to 30 µg of protein for the Triton fraction) were resolved on 4–12% SDS-PAGE gels (BioRad) and transferred to PVDF membranes using the Trans-Blot Turbo dry transfer system. Immunoblots were subsequently performed as described for PVDF membranes below.

For all other Western blots, cells were lysed in RIPA Buffer supplemented with protease and phosphatase inhibitors. The protein concentration of each sample was measured using Pierce BCA Protein Assay (Thermo Fisher). Samples were prepared with 20 µg of protein, 1x LDS sample buffer (Invitrogen NP0007) and 1x reducing agent (BioRad 1610972), and heated to 70°C for 10 minutes. Samples were loaded into 4-12% SDS-PAGE gels (BioRad), and a current of 120V was applied for approximately 60 minutes. The gel proteins were transferred to a PVDF membrane using BioRad TransBlot Turbo Transfer System, fixed in 4% PFA for 40 minutes and blocked in 5% w/v non-fat milk in 0.1% Tween in TBS (TBST) for 1 hour. For antibodies against phosphorylated proteins, blots were blocked in 5% w/v bovine serum albumin (BSA) in 0.1% TBST. Blots were then incubated with primary antibodies (dilutions specified in KRT) in blocking buffer overnight at 4°C. Membranes were washed in 0.1% TBST 3 times for 5 minutes each, incubated with secondary antibodies conjugated with horseradish peroxidase for 2 hours at room temperature in blocking buffer, and exposed to chemiluminescence activator before imaging using LI-COR Odyssey XF system. For dot blots, 3 µL of media was added onto nitrocellulose membranes and allowed to dry. Blocking and antibody incubation were performed as described for Western blots. Protein concentration in the media was quantified by Piece BCA Protein Assay and used to normalize dot blot signal.

### Collection and treatment with conditioned media

To generate neuron-conditioned media A53T-1A CORR28 neurons were plated at ∼150,000/cm2 on poly-L-ornithine and laminin-coated plates on day 7 of differentiation. The media was changed following the differentiation protocol described above. Starting day 14 of differentiation, the media was collected, centrifuged at 2000xg for 5 minutes to pellet debris, and stored at −20°C. Media was collected with each half media change, every 3-4 days.

To generate astrocyte-conditioned media, astrocytes were plated at ∼30,000/cm2 on 0.1% gelatin-coated plates in AM (ScienCell). The following day, media was changed to either fresh neuron media (Neurobasal, 1x B-27, 1x N-2, 1x MEM-NEAA, 1x GlutaMAX, 1% penicillin-streptomycin, 10ng/mL BDNF, 10ng/mL GDNF, 0.5 mM dcAMP, 1ug/mL laminin) supplemented with 10ng/mL CNTF or neuron conditioned media supplemented with 10ng/mL CNTF. After 3 days, media was collected, centrifuged at 2000xg for 5 minutes to pellet debris, and stored at −20°C.

To treat neurons with conditioned media, SNCA-A53T overexpressing neurons were plated at ∼150,000/cm2 on poly-L-ornithine and laminin-coated 96-well µClear plates (Greiner) on day 7 of differentiation, following the normal neuron differentiation protocol described above. On day 11 of differentiation, the media were changed to conditioned media supplemented with 5 µg/mL doxycycline and 1 µg/mL laminin. Half media changes with conditioned media supplemented with doxycycline and laminin were continued every 3-4 days until day 25 of differentiation, when cultures were fixed.

To treat microglia with conditioned media, SNCA-A53T neuron conditioned media was collected as described above and incubated with microglia cultured in 2D conditions after seeding at ∼85,000/cm2 on geltrex-coated 96-well µClear plates (Greiner).

### SNCA Knockdown Astrocyte Generation with shRNA Lentivirus

SNCA and scramble MISSION lentiviral shRNAs were purchased from Sigma Aldrich. SNCA shRNA lentivirus was produced as previously described^128^. Briefly, HEK293T cells were grown in Dulbecco’s Modified Eagle Medium (DMEM; Gibco 11965092) with 10% bovine calf serum (Cytiva SH30073) at 37 °C in a 5% CO_2_ incubator. The day before transfection, ∼10^7^ cells were plated per 15cm dish coated in 0.1% gelatin. Media was changed just prior to transfection. For one 15cm dish, 22.5ug of transfer vector was combined with 14.7ug of pMDLg/pRRE (Addgene plasmid #12251), 5.7ug of pRSV.Rev (Addgene plasmid #12253), and 7.9ug of pMD2.G (Addgene plasmid #12259) in a 15mL conical tube. The plasmid mixture was added to 1mL of 278 uM CaCl_2_ (Sigma C1016) and mixed thoroughly before adding 1mL of 2x BBS solution [280 mM NaCl (Fisher Scientific S271-1), 50 mM BES (Millipore 391334), 1.5mM Na_2_HPO_4_ (Sigma S5136); pH = 6.95] dropwise. Tubes were mixed by inversion, incubated at room temperature for 1 min, and then added dropwise to HEK293T dish. Cell culture media was changed after 16-24 h. Conditioned media was then collected every 24 hours for 2 days; conditioned media from the same plate was combined over the two collections. Conditioned media was centrifuged at 2000 x g for 10 minutes to pellet debris and filtered through 0.22-um-pore-size PES filter (Millipore SCGP00525). To concentrate virus, filtered media was incubated with PEG 8000 (Sigma P2139) and NaCl, at a final concentration of 5% and 0.15M respectively, overnight at 4°C, then centrifuged at 3000 x g for 20 minutes. The pellet was resuspended in DMEM/F12 with GlutaMAX (Gibco 10565018) to 1% of the original supernatant volume. The concentrated virus was stored in aliquots at −80°C. A detailed protocol for lentiviral production is available at doi.org/10.17504/protocols.io.6qpvr8ydolmk/v1.

iPSC-derived astrocytes were plated at ∼21,000 cells/um in a 6-well plate. The following day, cells were transduced at a 1:40 dilution of lentivirus in Astrocyte Medium (AM, ScienCell). Media was replaced with fresh AM 24 hours post transduction. Selection was started 72 hours post-transduction by adding fresh media supplemented with 2 ug/ml puromycin for two consecutive days. shRNA knockdown efficiency was validated by qPCR.

### Treatment with cholesterol-lowering drugs

Astrocytes were seeded into 0.1% gelatin-coated 6-well plates in AM (ScienCell). After 2-3 days, the media was changed to AM without FBS supplemented with cholesterol-lowering drugs (concentrations below) or DMSO as a vehicle. After 4 days, cells were seeded for final assays in 0.1% gelatin-coated 96-well µClear plates (Greiner) in AM with treatment. The following day, media was changed to FBS-free maturation media (50% DMEM/F12, 50% Neurobasal, 1X B27 without Vitamin A, 1X N2, 1X NEAA, 1X GlutaMAX, 10ng/mL CNTF, 1% penicillin-streptomycin) with treatment. Assays were started 2-3 days later, as described above. miBrains were treated with methyl-β-cyclodextrin (concentration below) or DMSO from day 0 of assembly to day 18, with half-media changes performed 3 times a week.

The following drug concentrations were used:

2-hydroxypropyl-β-cyclodextrin: 100 µM (Sigma C0926)
methyl-β-cyclodextrin: 100 µM (Cayman 21633)
atorvastatin: 50 nM (Sigma SML3030)
efavirenz: 10 µM (MedChemExpress HY10572)
LXR-623: 10 µM (MedChemExpress HY-10629)
T0901317: 10 µM (MedChemExpress HY-10626)

## QUANTIFICATION AND STATISTICAL ANALYSIS

All experimental data are presented as mean ± standard error of the mean. Statistical analyses were performed using GraphPad Prism (GraphPad Software). Data were assessed for normality using the D’Agostino-Pearson, Anderson-Darling, Shapiro-Wilk, and Kolmogorov-Smirnov tests to determine whether parametric or non-parametric statistical tests were used. Comparisons between two groups were performed using unpaired Student’s t-tests or Mann-Whitney tests, as appropriate. Comparisons among multiple groups were performed using one- or two-way ANOVA followed by appropriate multiple-comparisons tests. Statistical details, including the statistical test, sample size (n), and P values, for each experiment are provided in the results section and figure legends.

## ADDITIONAL RESOURCES

Detailed step-by-step protocols for all experimental procedures developed in this study are available in this study’s protocols.io collection: https://dx.doi.org/10.17504/protocols.io.n92ldomb9g5b/v1.

## Supplemental information

**Document S1.** Supplemental Figures S1-S5.

**Video S1. Neurons with α-synuclein pathological phenotypes within the miBrain**. 3D rendering with animated orthogonal projection of SNCA-A53T neurons within the miBrain. Green: SNCA-A53T-sfGFP; red: pS129 α-Syn; magenta: TUJ1, blue: Hoechst.

**Video S2. APOE3/3 astrocytes co-localize with neuronal-derived α-synuclein.** 3D rendering with animated orthogonal projection of astrocytes within an *APOE3/3* miBrain. Astrocytes display elongated morphology along neuronal networks. Green: SNCA-A53T-sfGFP; magenta: GFAP, blue: Hoechst.

**Video S3. *APOE4/4* reactive astrocytes poorly co-localize with neuronal-derived α-synuclein.** 3D rendering with animated orthogonal projection of astrocytes within an *APOE4/4* miBrain. Astrocytes display amoeboid morphology not aligned with neuronal networks. Green: SNCA-A53T-sfGFP; magenta: GFAP, blue: Hoechst.

**Table S1. Differentially expressed genes (DEGs).** Unfiltered DEGs in APOE3/3 A53T vs. Control and in APOE4/4 vs APOE3/3.

**Table S2. Pathway analysis.** Gene ontology biological processes enrichment and curated gene sets utilized for gene set enrichment analysis.

**Table S3. Metadata.** Cell-level metadata, including QC metrics and cell-type annotations.

**Table S4. QC.** QC metrics per sample.

**Table S5. Cell number.** Number of nuclei per sample and cellular cluster.

---

**Figure S1.**
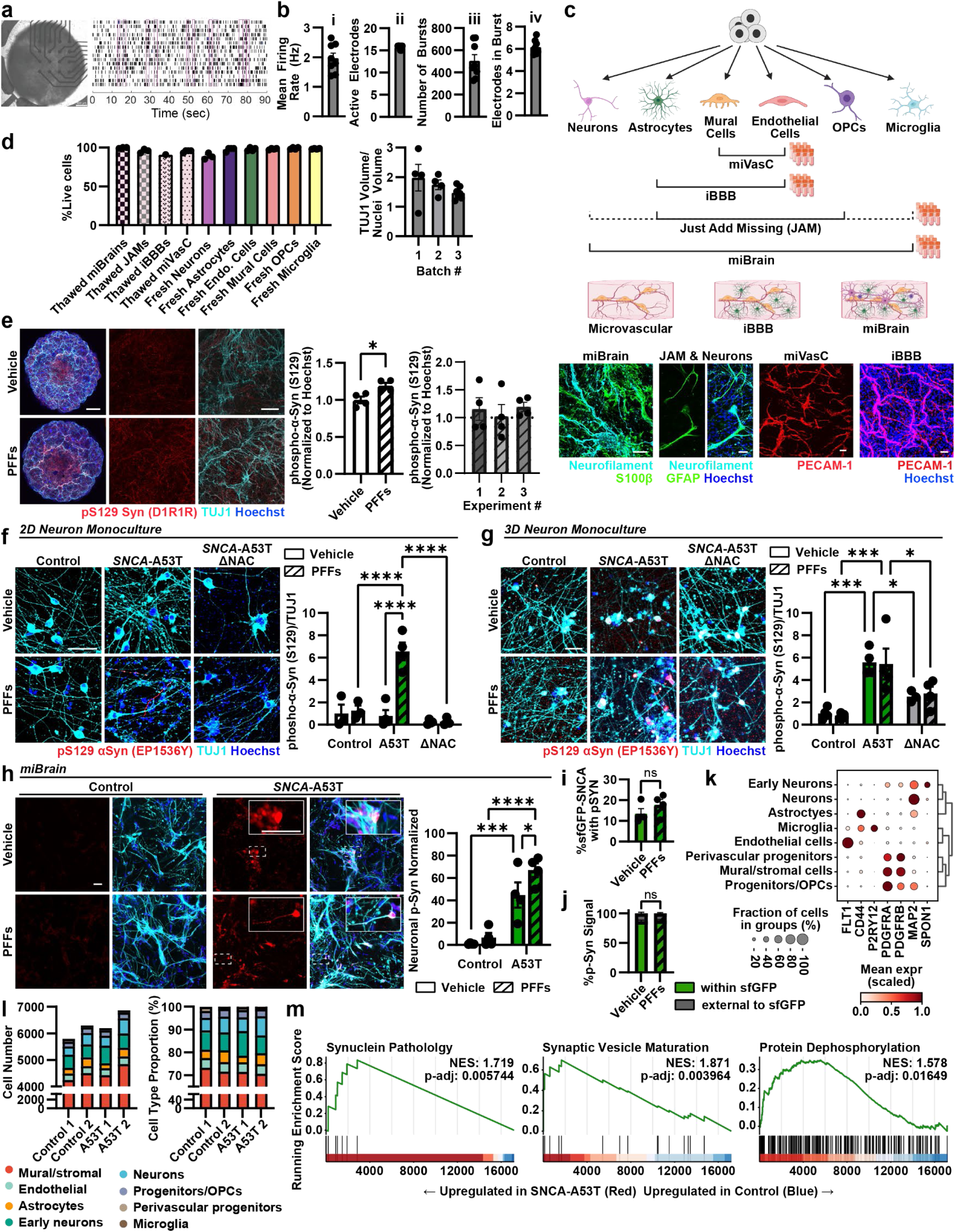
miBrain cryopreservation and development of α-synuclein intracellular inclusions, related to Figure 1. **a.** Representative microelectrode array (MEA) raster plot showing spontaneous neuronal firing across electrodes in 1-week old miBrains. Scale bar: 400 µm **b.** Quantification of MEA recordings: (i) mean firing rate (Hz), (ii) number of active electrodes per array, (iii) frequency of spontaneous bursts, and (iv) number of electrodes participating in synchronized network bursts. Data represents SEM (n = 8 biological replicates). **c.** Cartoon of the cryopreservation approach to preserve miBrains or smaller tissue units (miVasC: microvascular combo; BBB: blood-brain barrier; JAM: just add missing cell type). Representative images of two-week-old, thawed tissue stained with markers of neurons (neurofilament; cyan), astrocytes (S100b, GFAP; green), and endothelial cells (PECAM-1; red). Scale bars: 50 µm. **d.** Left: cell viability (% live) upon thawing the cryopreserved tissue compared with fresh cells harvested and counted. Data represent mean ± SEM (n = 1-4 biological replicates). Right: quantification of the ratio between neurons (TUJ1) and nuclei (Hoechst) volume in 18-day-old thawed miBrain tissue from three different batches. Microglia were not included in these frozen miBrains. Data represent mean ± SEM (n = 3 batches with 4-8 biological replicates each). **e.** Representative IF images of miBrains exposed to α-Syn pre-formed fibrils (PFFs) for 48h at week-2 in culture and stained after 14 days. Microglia were not included in these miBrains. α-Syn pS129 (red) intensity was normalized by nuclei (blue) and vehicle-treated miBrains; TUJ1 (cyan). Scale bars: 500 µm (low magnification) and 50 µm (high magnification). Data represent mean ± SEM (n = 4 biological replicates). P-values were calculated by a two-tailed, unpaired t-test. The left graph shows the quantification of the experiment represented by the images. The right graph shows the quantification across three different experiments (n = 4 biological replicates each). **f**. Representative IF images of α-Syn pS129 (red) in 25-DIV 2D-cultured neurons (TUJ1; cyan) generated from control (endogenous *SNCA*), *SNCA*-A53T, or *SNCA*-A53T-ΔNAC iPSCs, 14 days after vehicle or PFF treatment. α-Syn pS129 volume was normalized to TUJ1 volume and vehicle-treated control. Data represent mean ± SEM (n = 4 biological replicates). P-values were calculated by 2-way ANOVA and Tukey’s test. Scale bars: 50 µm. **g**. Representative IF images of α-Syn pS129 (red) in 25-DIV 3D-cultured neurons (TUJ1; cyan) generated from control (endogenous *SNCA*), *SNCA*-A53T, or *SNCA*-A53T-ΔNAC iPSCs, 14 days after vehicle or PFF treatment. α-Syn pS129 volume was normalized to TUJ1 volume and vehicle-treated control. Data represent mean ± SEM (n = 4 biological replicates). P-values were calculated by 2-way ANOVA and Tukey’s test. Scale bars: 50 µm. **h.** Representative IF images of α-Syn pS129 (red) in 18-days-old miBrains with control or *SNCA-*A53T neurons, 14 days after vehicle or PFF treatment. Microglia were not included in these miBrains. α-Syn pS129 volume within TUJ1 (cyan) was normalized to vehicle-treated control miBrains. Data represent mean values ± SEM (n = 6 biological replicates). P-values were calculated by 2-way ANOVA and Fisher’s LSD. **i.** Quantification of % area of sfGFP-SNCA co-stained with α-Syn pS129. Data represent mean values ± SEM (n = 4 biological replicates). P-values were calculated by a two-tailed, unpaired t-test. **j.** Quantification of α-Syn pS129 co-localized with sfGFP or external to sfGFP. Data represent mean values ± SEM (n = 4 biological replicates). P-values were calculated by 2-way ANOVA and Fisher’s LSD. *p < 0.05, **p < 0.01, ***p < 0.001, ****p < 0.0001. **k.** Dot plot of the expression levels and percentage of nuclei expressing specific cell markers across all annotated cell types in all miBrains (control, *APOE3/3* A53T and *APOE4/4* A53T). **l.** Number of cells n of each cell type (left) and proportion of each cell type (right) in control and *SNCA-*A53T miBrains. **m.** Representative gene set enrichment analysis (GSEA) plots showing α-synuclein pathology, synaptic maturation and protein dephosphorylation are modulated in *SNCA*-A53T miBrains compared to control.

**Figure S2:**
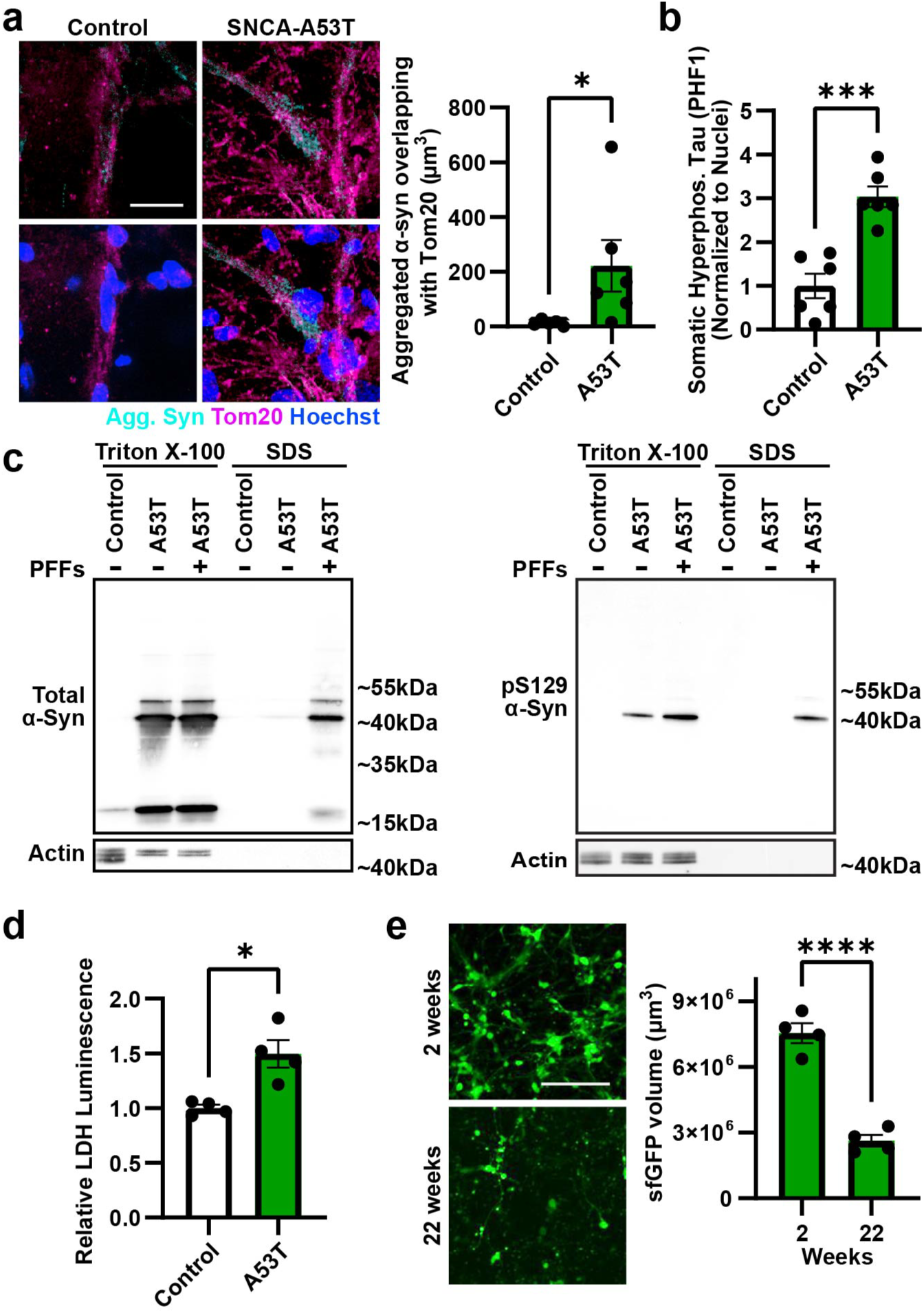
Characterization of α-Syn inclusions in miBrain tissue, related to Figure 2. **a.** Representative IF images of control and *SNCA*-A53T 18-days-old miBrains stained for aggregated α-Syn (cyan) and TOM20 (magenta). Volume of co-localized aggregated α-Syn and TOM20 was normalized to control miBrains. Data represent mean ± SEM (n = 6 biological replicates). P-value was calculated by the Mann-Whitney test. Scale bar: 25 µm. **b.** Quantification of total hyperphosphorylated tau (PHF1) in control and *SNCA*-A53T miBrains. Representative images in Figure 2b. PHF1 volume normalized to the number of nuclei and control miBrains. Data represent mean ± SEM (n = 6 biological replicates). P-value was calculated by a two-tailed, unpaired test. **c.** Western blots of total α-Syn (left) and α-Syn pS129 (right). Protein was extracted sequentially in Triton X-100 and SDS lysis buffers from control or *SNCA-*A53T neurons grown in 2D monoculture, with or without PFF treatment. Actin was used as a loading control. **d.** Quantification of relative lactate dehydrogenase (LDH) levels in control and *SNCA*-A53T miBrain media after 14 days of culture. Bars represent mean ± SEM (n = 4 biological replicates). P-value was calculated by Welch’s two-tailed t-test. **e.** Representative images of SNCA-A53T-sfGFP in live miBrains after 2 and 22 weeks of culture. Data represent mean sfGFP volume ± SEM (n = 4 biological replicates). P-value was calculated by a two-tailed, unpaired t-test. Scale bar: 100 µm. *p < 0.05, **p < 0.01, ***p < 0.001, ****p < 0.0001

**Figure S3.**
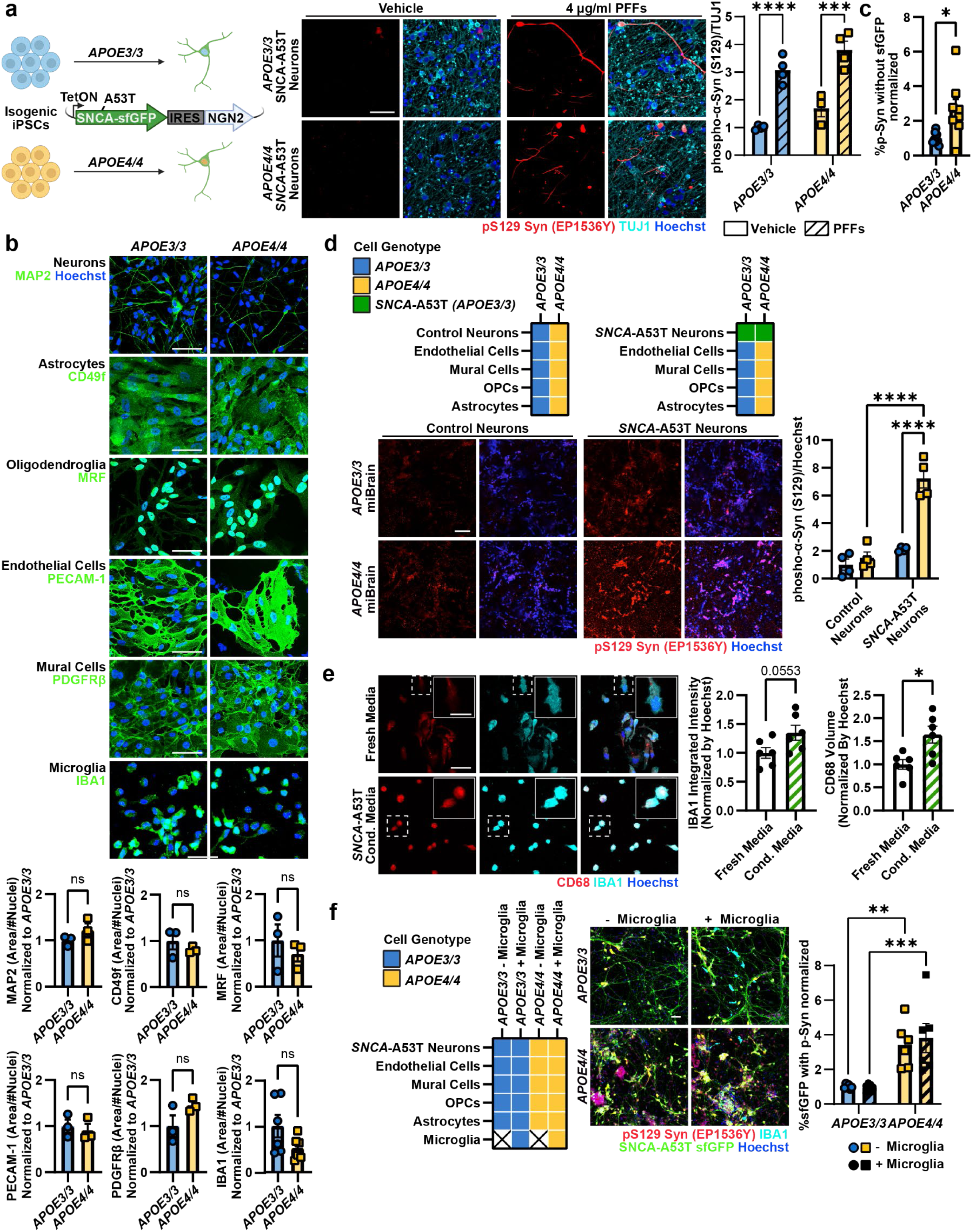
Non-neuronal cells promote α-synuclein pathological phenotypes in *APOE4/4* tissue, related to Figure 3. **a.** Representative IF images of α-Syn pS129 (red) in 2D-cultured *APOE3/3* and *APOE4/4* isogenic 25-DIV neurons overexpressing SNCA-A53T-sfGFP, 14 days after vehicle or PFF treatment. α-Syn pS129 volume was normalized to TUJ1 (cyan) volume and *APOE3/3* vehicle treated cells. Data represent mean ± SEM (n = 4 biological replicates). P-values were calculated by 2-way ANOVA and Fisher’s LSD. Scale bar: 50 mm. **b.** Representative IF images of monocultures of all cell types of the miBrain, stained for cell-specific markers (green). Positive cell area was normalized to the number of nuclei (blue) and *APOE3/3* cells. Data represent mean ± SEM (n = 3 biological replicates). P-values were calculated by two-tailed, unpaired t-tests. Scale bar: 50 µm. **c.** Quantification of α-Syn pS129 outside of sfGFP volume (non-neuronal). Normalized to *APOE3/3* miBrains. Data represent mean ± SEM (n = 8 biological replicates). P-value was calculated by Welch’s two-tailed t-test. **d.** Representative IF images of α-Syn pS129 (red) in *APOE3/3* and *APOE4/4* 18-days-old miBrains with control or *SNCA*-A53T neurons. The A53T neurons were generated from the A53T-1A CORR28 line (donor with familial Parkinson’s disease). α-Syn pS129 volume was normalized to Hoechst (blue) volume and *APOE3/3* miBrains with control neurons. Data represent mean ± SEM (n = 4 biological replicates). P-values were calculated by 2way ANOVA and Fisher’s LSD. Scale bar: 50 µm. **e.** Representative IF images of IBA1 (cyan) and CD68 (red) in 2D-microglia monoculture after exposure to fresh or SNCA-A53T neuron conditioned media. IBA1 intensity and CD68 volume were normalized by Hoechst volume (blue) and cells exposed to fresh media conditions. Data represent mean ± SEM (n = 6 biological replicates). P-values were calculated by two-tailed, unpaired t-tests. Scale bar: 50 µm (low magnification) and 25 µm (high magnification insets). **f.** Representative IF images of α-Syn pS129 (red) in *APOE3/3* and *APOE4/4* 18-days-old miBrains, with or without isogenic microglia (IBA1; cyan). α-Syn pS129 volume within SNCA-A53T-sfGFP (green) neurons was normalized to sfGFP volume and *APOE3/3* miBrains without microglia. Data represent mean ± SEM (n = 6 biological replicates). P-values were calculated by 2way ANOVA and Fisher’s LSD. Scale bar: 50 µm. *p < 0.05, **p < 0.01, ***p < 0.001, ****p < 0.0001.

**Figure S4.**
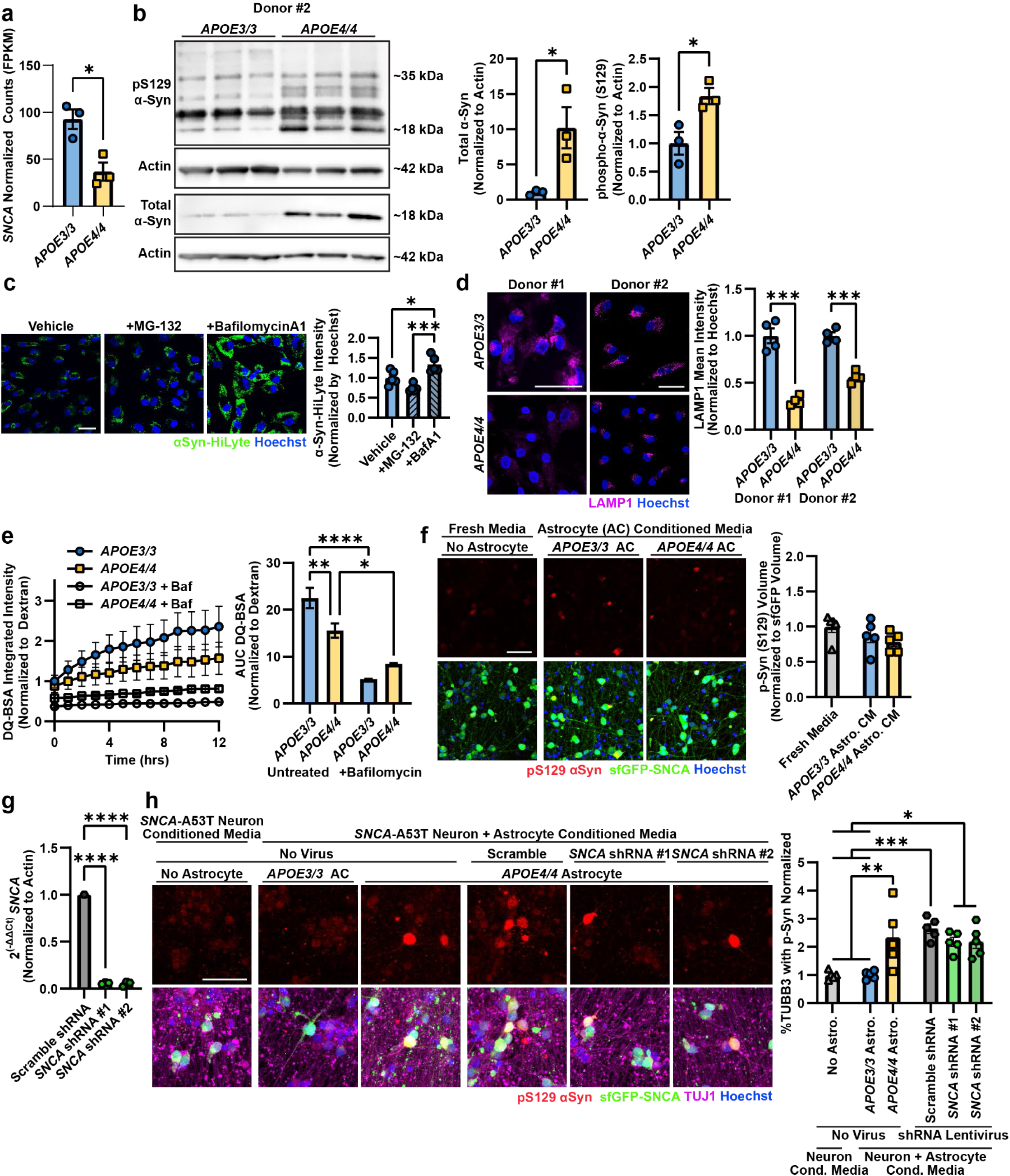
Characterization of lysosomal function and α-Synuclein uptake and accumulation in astrocytes, related to Figure 4. **a.** RNAseq analysis of isogenic, iPSC-derived astrocytes for SNCA expression using normalized counts (FPKM). Data represent mean ± SEM (n = 3 biological replicates). P-value was calculated by a two-tailed, unpaired t-test. **b.** Western blots of *APOE3/3* and *APOE4/4* astrocytes for total α-Syn protein and phosphorylated α-Syn protein. β-Actin was used as a loading control. Quantifications normalized to β-Actin and *APOE3/3.* Data represent mean ± SEM (n = 3 replicates). P-values were calculated by Welch’s one-tailed t-test (total α-Syn) and unpaired, one-tailed t-test (pS129 α-Syn) **c.** Representative IF images of astrocytes with α-Syn-HiLyte (green) after 24 hours of uptake and 24 hours of degradation after treatment with BafilomycinA1 and MG-132. αSyn-HiLyte intensity was normalized to Hoechst (blue) area and vehicle-treated astrocytes. Data represent mean ± SEM (n = 5 biological replicates). P-values were calculated by one-way ANOVA and Sidak’s test. Scale bar = 50 µm. **d.** Representative IF images of LAMP1 immunostaining (magenta) in two different isogenic iPSC lines. LAMP1 intensity normalized to Hoechst (blue) and *APOE3/3* for each isogenic pair. Data represent mean ± SEM (n = 4 biological replicates). P-values were calculated by two-tailed, unpaired t-tests. Scale bar = 50 µm. **e.** DQ-BSA Red integrated intensity normalized to dextran and measured over 24 hours on an Incucyte (Sartorius). 100 nM bafilomycin A1 was used as a control for non-lysosomal proteolysis of DQ-BSA. Data represents mean ± SEM (n = Data points represent mean values and error bars represent standard error (n = 3-5 biological replicates). P-values were calculated by 2-way ANOVA and Fisher’s LSD on area under the curve values. **f.** Representative IF images of pS129 α-Syn (red) in *SNCA*-A53T neurons treated with conditioned media from *APOE3/3* or *APOE4/4* astrocytes or with fresh neuron media. pS129 α-Syn volume normalized to SNCA-A53T-sfGFP (green) volume and to cells treated with fresh media. Data represent mean ± SEM (n = 6 biological replicates). P-values were calculated by one-way ANOVA and Tukey’s test. Scale bar = 50 µm. **g**. qPCR validation of lentiviral shRNAs targeting *SNCA* in astrocytes. Relative gene expression was calculated by ΔΔCt with *ACTB* as a housekeeping gene. Data represent mean ± SEM (n = 1-3 biological replicates). P-values were calculated by one-way ANOVA and Dunnett’s test. **h.** Representative IF images of pS129 α-Syn (red) in *SNCA*-A53T neurons treated with neuron-conditioned media or neuron and astrocyte-conditioned media from *APOE3/3, APOE4/4,* or *APOE4/4* with *SNCA* shRNA astrocytes. pS129 α-Syn volume was normalized to TUJ1 (magenta) volume and neuron-conditioned media samples. Data represent mean ± SEM (n = 5 biological samples). P-values were calculated by one-way ANOVA and Tukey’s test. Scale bar = 50 µm. *p < 0.05, **p < 0.01, ***p < 0.001, ****p < 0.0001.

**Figure S5.**
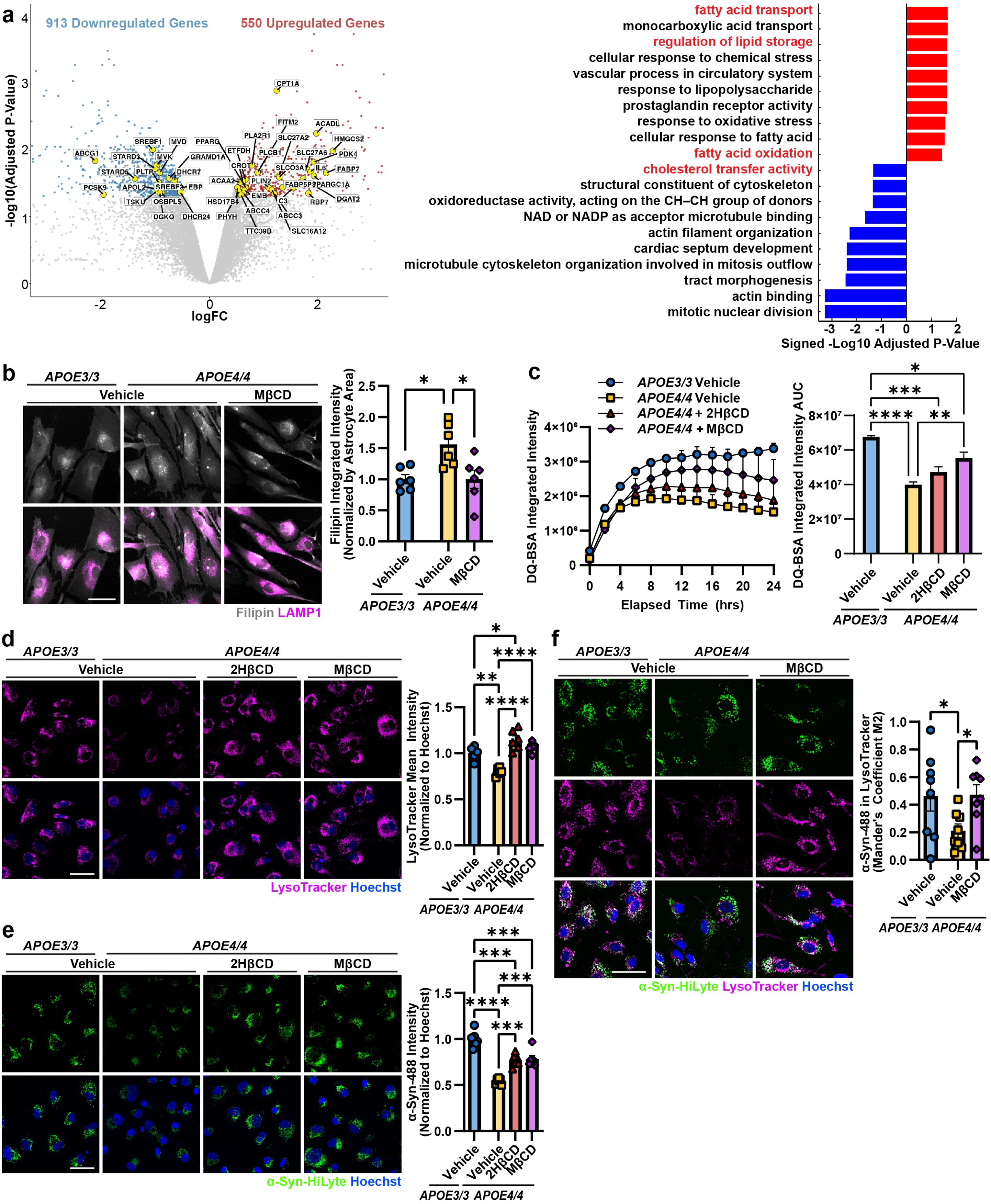
MβCD treatment in *APOE4/4* astrocytes improves lysosomal uptake of α-Synuclein, related to Figure 5. **a.** Volcano plot of *APOE4/4* vs. *APOE3/3* astrocytes showing DEGs (p adj < 0.5, abs(logFC > 0.5). Top 10 Gene Ontology (GO) upregulated (red) and downregulated (blue) pathways ranked by adjusted p-value. Genes highlighted in the volcano plot correspond to GO terms related to lipid metabolism b. Representative IF images of Filipin (grey) and LAMP1 (magenta) in astrocytes treated with MβCD. Filipin integrated intensity was normalized by astrocyte area and *APOE3/3.* Data represent mean ± SEM (n = 6 biological replicates). P-values were calculated by 2-way ANOVA and Tukey’s test. Scale bar = 50 µm. **c.** Left: DQ-BSA Red integrated intensity in astrocytes with 2HβCD or MβCD treatment, measured over 24 hours on an Incucyte (Sartorius) in a second isogenic donor line (Donor #3). Data represent mean ± SEM (n = 4 biological replicates). Right: Area under the curve calculation for DQ-BSA. Data represent mean ± SEM (n = 4 biological replicates). P-values were calculated by 2-way ANOVA and Tukey’s test. **d.** Representative IF images of LysoTracker (magenta) in astrocytes treated with cyclodextrins in a second isogenic donor line (Donor #3). LysoTracker intensity was normalized by Hoechst (blue) area and *APOE3/3*. Data represent mean ± SEM (n = 5-6 biological replicates). P-values were calculated by 2-way ANOVA and Tukey’s test. Scale bar = 50 µm. **e.** Representative IF images of astrocytes treated with cyclodextrins after a 24h incubation with fluorescently labeled α-Syn-HiLyte (green) in a second isogenic donor line (Donor #3). α-Syn-HiLyte intensity was normalized by Hoechst (blue) area and *APOE3/3*. Data represent mean ± SEM (n = 6 biological replicates). P-values were calculated by 2-way ANOVA and Tukey’s test. Scale bar = 50 µm. **f**. Representative IF images of fluorescently labeled α-Syn-HiLyte (green) co-localized with LysoTracker (magenta) in astrocytes treated with cyclodextrins. Calculated Mander’s coefficient M2 (amount of α-Syn signal overlapping with LysoTracker signal). Data represent mean ± SEM (n = 8 replicates). P-values were calculated by 2-way ANOVA and Fisher’s test. Scale bar = 50 µm. *p < 0.05, **p < 0.01, ***p < 0.001, ****p < 0.0001.

